# Cell targeting and immunostimulatory properties of a novel Fcγ-receptor independent agonistic anti-CD40 antibody in rhesus macaques

**DOI:** 10.1101/2023.03.22.533762

**Authors:** Xianglei Yan, Sebastian Ols, Rodrigo Arcoverde Cerveira, Klara Lenart, Fredrika Hellgren, Kewei Ye, Alberto Cagigi, Marcus Buggert, Falk Nimmerjahn, Jesper Falkesgaard Højen, Daniel Parera, Ulrich Pessara, Stephan Fischer, Karin Loré

## Abstract

Targeting CD40 by agonistic antibodies used as vaccine adjuvants or for cancer immunotherapy is a strategy to stimulate immune responses. The majority of studied agonistic anti-human CD40 antibodies require crosslinking of their Fc region to inhibitory FcγRIIb to induce immune stimulation although this has been associated with toxicity in previous studies. Here we introduce an agonistic anti-human CD40 monoclonal IgG1 antibody (MAB273) unique in its specificity to the CD40L binding site of CD40 but devoid of Fcγ-receptor binding, we demonstrate rapid binding of MAB273 to B cells and dendritic cells resulting in strong activation *in vitro* on human cells and *in vivo* in rhesus macaques. Dissemination of fluorescently labeled MAB273 after subcutaneous administration was found predominantly at the site of injection and specific draining lymph nodes. Phenotypic cell differentiation and upregulation of genes associated with immune activation were found in the targeted tissues. Antigen-specific T cell responses were enhanced by MAB273 when given in a prime-boost regimen and for boosting low preexisting responses. MAB273 may therefore be a promising immunostimulatory adjuvant that warrants future testing for therapeutic and prophylactic vaccination strategies.

## Introduction

In the past decade, development of targeted therapies that block immune checkpoint inhibitors has revolutionized immunotherapy. However, blocking immune checkpoints alone such as programmed cell death protein 1 (PD-1) or cytotoxic T-lymphocyte-associated protein 4 (CTLA-4) is not sufficient as most patients do not induce long-term sustained and effective responses[1–3]. Additional strategies like the CD40/CD40L interaction has been explored to specifically target immune cells to enhance antigen presentation and immunity but has also shown variable success[4]. CD40 is a member of the tumor necrosis factor receptor superfamily (TNFRSF) predominantly expressed on the cell-surface of antigen presenting cells (APCs) including B cells, dendritic cells (DCs) and monocytes and works as a costimulatory receptor[5]. CD40 binds to CD40 ligand (CD40L, CD154) on T cells during the antigen presentation process which leads to strong activation of both APCs and T cells[6]. Previous studies have shown that CD40 activation can replace T cell help required to drive CD8 T cell responses[7–9]. CD40 agonists have therefore been of interest to develop as candidates for adjuvants or cancer immunotherapy[10].

Among the CD40 agonists that have progressed to clinical development, CP-870,893 (Selicrelumab) is the most advanced that has been tested for treatment of several solid tumors, especially for melanoma[11–14]. While it has been shown that a single dose can induce antitumor activity in some but not all of the individuals, the main adverse events were cytokine release syndrome (CRS) with grade 1 to 2, transient liver function test abnormalities and transient decrease in platelet counts[11–13, 15]. More information is needed of the mechanisms by which CD40 agonists can mediate beneficial immune stimulation without unwanted side effects in order to refine CD40 agonists further. A proposed strategy to improve safety is to modify the fragment crystallizable region (Fc region) of CD40 antibodies.

Among the Fc-gamma receptors (FcγRs), binding to FcγRIIb has been shown to potently enhance the activity of CD40 agonists[16–18]. The new generation of CD40 agonists with an engineered Fc region to increase binding to FcγRIIb has therefore been developed[18–22]. However, by introducing mutations in CP-870,893 to increase the affinity to FcγRIIb side effects were increased[20, 21]. These results suggest that in addition to the canonical CD40 signaling leading to antigen-specific T cell licensing, alternative pathways such as FcγRIIb crosslinking may also induce adverse events that are difficult to control. Manipulating or reducing FcγRIIb or even FcγR crosslinking in general is one strategy to develop safe but potent CD40 agonistic antibodies.

MAB273 is a novel humanized rabbit IgG1 with the LALA (L234A and L235A)-mutations in the Fc region which disables Fcγ-receptor-mediated crosslinking[23]. In this study, we tested the CD40 binding and immune activation capacity of MAB273 in human and rhesus macaque peripheral blood mononuclear cells (PBMCs) and confirmed that this was independent of the Fc region. The tolerability and immunostimulatory functions of MAB273 *in vivo* were further tested in rhesus macaques by analyzing biodistribution in different tissues and cells, innate immune activation and induction of antigen-specific T cells.

## Materials and Methods

### Antibodies and the generation of fragments

MAB273, CP-870,893 and isotype control antibody IgG1-LALA were provided by Icano MAB GmbH. For generation of F(ab’)2 and Fab fragments of MAB273, after pepsin (for F(ab’)2) or papain (for Fab) digestion, a CH1-XP column (ThermoFisher, A37054) containing anti-IgG-CH1 matrix which binds to the CH1 region of IgG was used, thereby only undigested MAB273 and F(ab’)2 or Fab fragments remained after positive selection. A Fc-XP column (ThermoFisher, 494371201) containing anti-IgG-CH3 matrix which binds to the CH3 region (on Fc region) of IgG was used to process sample from last step, the undigested MAB273 which still has Fc region was removed and only F(ab’)2 or Fab fragments were remained by negative selection.

### Human samples

The work on human blood samples was approved by the Swedish Institutional Review Board of Ethics. The informed consent was signed according to the Declaration of Helsinki. All blood samples were not associated with the features that could be linked to identification (such as sex and age). Sample size is indicated in each figure legend.

### Animals

This study was approved by the Local Ethical Committee on Animal Experiments. Six male Indian-origin rhesus macaques (for toxicity and safety study) and three male and three female Indian-origin rhesus macaques (for immunogenicity and biodistribution study) were housed at Astrid Fagraeus Laboratory, Karolinska Institutet. All procedures were performed according to the guidelines of the Association for Assessment and Accreditation of Laboratory Animal Care.

### Immunizations and sampling

In the initial toxicity and safety study, the animals were divided into three groups. Two animals received intravenous (i.v.) administration of 1 mg/kg anti-CD40 mAb (MAB273), two animals received 0.1 mg/kg MAB273 by i.v., and the last two animals received subcutaneous (s.c.) administration of 0.1 mg/kg MAB273. For i.v. administration, an intravenous infusion of the antibody into the saphenous vein was performed, a total volumeof 25ml was slowly distributed stepwise for 30 minutes. For s.c. administration, a single 0.5ml subcutaneous injection of the antibody in the skin above the left quad muscle was performed. Blood draws were performed at pre-dose, 30 minutes, 4 hours, 24 hours, 48 hours, 72 hours, 1 week, 2 weeks, 3 weeks, and 4 weeks after MAB273 administration in heparin tubes.

In the immunogenicity study, animals were divided into two groups. In the therapeutic vaccination group, animals were immunized s.c. with 0.1 mg/kg Env peptides[24] (Pep1|YLRDQQLLGIWG, Pep2|RQQQNNLLRAIEA, Pep3|VYYGVPVWKEA, Pep4|LWDQSLKPCVKLT, Pep5|SVITQACSKVSFE, Pep6|GTGPCTNVSTVQC, Pep7|YKVVKIEPL, GenScript, New Jersey, U.S.) to establish low immunity during prime and boost (before 11 weeks). They thereafter received 1 mg/kg Env peptides plus 0.1 mg/kg MAB273 s.c. to measure the enhancement effect after exposure to the second boost (at 11 weeks). In the prophylactic vaccination group, animals were co-injected with 0.1 mg/kg Env peptides and 0.1 mg/kg MAB273 s.c. after prime and boost (at 7 weeks) and with 1 mg/kg Env peptides and 0.1 mg/kg MAB273 s.c. during the second boost (at 11 weeks) and with 1 mg/kg Env peptides, 0.1 mg/kg MAB273 and 100 μg Env protein (426c NFL trimer, provided by Richard Wyatt, Scripps Research) during the third boost (at 24 weeks). Blood draws were performed at pre-dose, 48 hours, 1 week, 2 weeks, 3 weeks, 7 weeks, 8 weeks, 9 weeks, 11 weeks (48 hours after the 2^nd^ boost time point was also sampled), 12 weeks, 13 weeks after the first administration in heparin tubes, the prophylactic vaccination group was followed for additional 4 weeks. Bronchoalveolar lavage (BAL) sampling was performed as described previously[25] at 9 weeks and 13 weeks after the first administration.

For biodistribution assessment, three animals were immunized with 0.1 mg/kg Alexa Fluor 680-labeled MAB273 and terminated after 24 (one animal) or 48 hours (two animals). Multiple samples were collected, such as skin biopsies from the site that received MAB273, the skin site on the contralateral leg that received saline, draining left and right inguinal LNs, left and right common iliac LNs, paraaortic LNs, mediastinal LNs, PBMCs, bone marrow, spleen, liver and BAL. The samples were processed into single-cell suspensions as described previously[26–28].

### Safety clinical chemistry and hematology tests

Hematological analyses of heparinized blood were performed within 8 h after collection on an Exigo Vet instrument (Model H400, Boule Diagnostics AB, Spånga, Sweden) after QC with use of Boule Vet Con control blood. Heparinized plasma samples were analyzed using an ABAXIS Vetscan VS2 3.1.35 Chemistry analyzer (Triolab, Solna, Sweden). Indicated parameters were analyzed on Mammalian Liver Profile rotors (Triolab), which have individual QC controls.

### Blood sample processing

PBMCs were isolated by Ficoll-Paque (GE Healthcare, Fairfield, CT) density gradient centrifugation of blood samples at 2200 rpm for 25 minutes with no brake or acceleration. PBMCs were washed and maintained in phosphate-buffered saline (PBS). Samples were stained immediately or frozen using 90% heat-inactivated fetal bovine serum (FBS) and 10% DMSO (Sigma-Aldrich) and stored at −170°C.

### Flow cytometry for innate phenotyping

In the *in vitro* assays, fresh PBMCs were exposed to antibodies for 2 hours at 4°C (CD40 binding) or 24 hours at 37°C (activation) first. In the *in vivo* assays, fresh PBMCs were stained with LIVE/DEAD™ Fixable Blue Dead Cell Stain Kit (Invitrogen, L23105), then blocked with FcR Blocking Reagent (Miltenyi Biotec, 130-059-901) according to manufacturer’s protocol. Samples were then surfaced stained with a panel of fluorescent staining antibodies (Table S1). After staining and washing, PBMCs were resuspended in 1% paraformaldehyde (PFA) and acquired on an LSRFortessa flow cytometer (Fortessa, BD). Data analysis was performed using FlowJo v10.

### Flow cytometry for labeled antibody tracking

Single-cell suspensions from collected samples were stained with LIVE/DEAD then blocked with FcR Blocking. Cells were then surfaced stained with a panel of fluorescent staining antibodies (Table S1). After staining and washing, cells were spiked with AccuCount beads (Spherotech, ACBP-100-10) and resuspended in 1% PFA and acquired on an LSRFortessa flow cytometer. Counting bead normalized cell numbers were calculated according to the manufacturer’s protocol.

### Flow cytometry for detecting antigen-specific T cells

Fresh PBMCs were stimulated with 2 μg/mL Env peptides or 10 μg/mL Env protein overnight at 37°C, BV421-CD107a staining antibody (BioLegend, 328626) was added during the incubation. On the next day, GolgiStop (Monensin, BD, 554724) and Golgi Plug (Brefeldin A, BD, 555029) were added 6 hours before staining according to manufacturer’s protocol. The LIVE/DEAD staining was the same as above, samples were then surfaced stained, permeabilized with Cytofix/Cytoperm™ (BD, 554714), and intracellular staining performed (Table S1). After staining and washing, PBMCs were resuspended in 1% PFA and acquired on an LSRFortessa flow cytometer, the background was subtracted with unstimulated autologous controls.

### Flow cytometry for B cell proliferation

Fresh PBMCs were labeled with 0.25µM CellTrace Violet (Invitrogen) at a cell concentration of 1 million/mL for 20 min at 37°C. Labeled PBMCs were stimulated with testing Abs. As controls, cells were stimulated with either 1µg/ml CpG B (Invivogen) or left unstimulated in complete media (RPMI1640, 10% FBS, 1% L-glutamine, 1% penicillin/streptomycin) and cultured for 5 days. After culture, cells were washed with PBS and stained with LIVE/DEAD and then blocked with FcR Blocking. Samples were then surfaced stained with a panel of fluorescent staining antibodies (Table S1). After staining and washing, PBMCs were resuspended in 1% PFA and acquired on an LSRFortessa flow cytometer.

### Flow cytometry for CD40L competition

Fresh PBMCs were cultured with different concentration of MAB273, CP-870,893 or CD40L (ThermoFisher, 34-8902-81) for 20 minutes at 4°C then washed with cold PBS, 1 μg/mL CD40L-biotin (ThermoFisher, 15836427) was added for 20 minutes at 4°C then washed with cold PBS. The LIVE/DEAD staining was the same as above, samples were then surfaced stained with a panel of fluorescent staining antibodies (Table S1). After staining and washing, PBMCs were resuspended in 1% PFA and acquired on an LSRFortessa flow cytometer.

### ELISA assay for CD40L competition

Greiner-Bio One 96 well half-area ELISA plates (VWR, 738-0032) were coated overnight at 4°C with 2 µg/ml CD40 protein (ThermoFisher, A42565) in fresh PBS. The plates were blocked with PBS containing 5% (w/v) milk for 1 hour at room temperature (RT). Serially diluted MAB273, CP-870,893, or CD40L were added to plates and incubated for 2 hours at RT. Then, 2 μg/mL CD40L-biotin was added to plates and incubated for 1 hours at RT. The CD40L-biotin was detected by adding a 1:1,000 dilution of Streptavidin-HRP (Mabtech, 3310-9-1000) and the signal was developed by addition of TMB substrate (BioLegend). The addition of an equal volume of 1M H_2_SO_4_ stopped the reaction, and the optical density (OD) was read at 450 nm and background was read at 550 nm. The plates were washed 3 times between each incubation step using PBS supplemented with 0.05% Tween 20.

### ELISA assay for CD40 binding

ELISA plates were coated with 2 µg/ml CD40 protein and blocked as described above. Serially diluted MAB273, its Fab or F(ab’)2 fragments were added to plates and incubated for 2 hours at RT. The CD40 binding signal was detected by adding a 1:5,000 dilution of goat anti-human Fab/F(ab’)2 IgG secondary-HRP antibody (Jackson ImmunoResearch, 109-035-006) or a 1:20,000 dilution of goat anti-human Fc IgG secondary-HRP antibody (Jackson ImmunoResearch, 109-035-008) and the signal was developed as described above.

### MAB273 fluorochrome-labeling

MAB273 was labeled by using the Alexa Fluor™ 680 Protein Labeling Kit (ThermoFisher, A20172) according to manufacturer’s protocols. The Alexa Fluor 680-conjugated MAB273 was assessed for signal intensity and activation capacity by staining PBMCs overnight then acquire PBMCs on an LSRFortessa flow cytometer. The capacity to bind CD40 was tested by ELISA and compared to non-conjugated MAB273 as described above.

### ELISA assay for detection of anti-MAB273 IgG in plasma

ELISA plates were coated with 1 µg/ml MAB273 and blocked as described above. Serially diluted plasma samples were added to plates and incubated for 2 hours at RT. The anti-MAB273 IgG was detected by adding a 1:5,000 dilution of anti-macaque pan-species IgG HRP antibody (Absolute Antibody, clone 1B3) and the signal was developed as described above.

### ELISA assay for detection of anti-Env peptides IgG

ELISA plates were coated with 2 µg/ml NeutrAvidin (ThermoFisher, 31000) and blocked as described above. Then, 2 µg/mL biotin conjugated Env peptides (GenScript, customized) were added to plates and incubated for 1 hour at RT. Serially diluted plasma samples were added to plates and incubated for 2 hours at RT. The anti-Env peptides IgG was detected by adding a 1:5,000 dilution of polyclonal anti-monkey IgG HRP antibody (Nordic MUBio, GAMon/IgG(Fc)/PO) and the signal was developed as described above.

### ELISA assay for detection of anti-Env protein IgG

ELISA plates were coated with 2 µg/ml mouse anti-His tag antibody (R&D Systems, MAB050) and blocked as described above. Then, 2 µg/mL Env protein (NFL 426c, His-tag) was added to plates and incubated for 1 hour at RT. Serially diluted plasma samples were added to plates and incubated for 2 hours at RT. The remaining steps are the same as above.

### ELISA assay for MAB273 pharmacokinetics analysis

ELISA plates were coated with 2 µg/ml CD40 protein and blocked as described above. Serially diluted plasma was added to plates and incubated for 2 hours at RT. MAB273 was detected by adding a 1:5,000 dilution of monkey cross-adsorbed polyclonal goat anti-human IgG HRP antibody (Southern Biotech, 2049-05) and the signal was developed as described above.

### ELISA assays for detection of cytokines

Supernatant from rhesus cell culture or rhesus plasma samples were evaluated for IL-12 p40, IL-6, IFN-γ and TNF levels by ELISA kits (Mabtech, 3450-1H-6, 3460-1H-6, 3421M-1H-6, 3512M-1H-6). Assays were performed according to manufacturer’s protocols.

### Bulk transcriptomics

Skin punch biopsy (MAB273 injection site and saline injection site), inguinal lymph nodes (MAB273 injection site and saline injection site), and blood (pre-immunization and 24-48 hours after immunization) samples were collected and preserved in RNAlater™ Stabilization Solution (ThermoFisher, AM7021) or PAXgene® Blood RNA Tube (BD, 762165). RNA isolation, library preparation, and sequencing were processed at the BEA core facility, Karolinska Institutet, using Illumina stranded mRNA prep kit. Illumina NovaSeq 6000 platform was used to generate paired-end reads of 150 bp with an average sequencing depth of 40 million reads per sample. Samples were preprocessed using nf-core rnaseq pipeline (version 3.7), genome alignment was processed using STAR alignment (version 2.7.10a) to the *Macaca mulatta* genome (Mmul_10) and quantification with Salmon (version 1.8.0).

### RNA sequencing data analysis

For this study, we used a customized bioinformatic analysis workflow of RNA sequencing data using R (version 4.1.2). Differential gene expression analysis was performed using DESeq2 (version 1.34.0). Gene Set Enrichment analysis was done with ClusterProfiler (version 4.2.2) package. The database used for gene set enrichment analysis was the Blood Transcriptome Modules (BTMs)[29]. To compare differentially expressed genes, Wald test was performed with multiple hypothesis testing controlling the false discovery rate (FDR) using the Benjamini-Hochberg procedure (*q-value* < 0.05).

### Statistics

No statistical methods were used to predetermine sample size. A Wilcoxon matched-pairs signed-rank test was used when two groups were compared and a Friedman test was used when three or more groups were compared, the results were considered statistically significant when p < 0.05, indicated as *p < 0.05 in the figures. Analyses were performed in GraphPad Prism 9.

## Results

### MAB273 binds the CD40L binding site and activates immune cells

Screening and identification of MAB273 have recently been described[23]. In this study, we focused on further functional characterization of MAB273. Since CP-870,893 is one of the most well-studied and potent CD40 antibodies in clinical development we used it as a comparator in the *in vitro* assays. CD40 binding capacity and activation were analyzed on multiple immune cells but we focused on B cells and myeloid dendritic cells (MDCs) within the PBMC population since they are central in immunity and express high levels of CD40 (Figure S1A). Human PBMCs exposed to MAB273 or CP-870,893 were found to have markedly reduced signal of a CD40 staining antibody (clone: 5C3) confirming that they both compete for binding to CD40 in a dose dependent manner (Figure 1A). However, using a CD40L competition assay only MAB273 binding was affected demonstrating that the epitope specificity on CD40 is different between MAB273 and CP-870,893 and only MAB273 binds the CD40L binding site (Figure 1B). Regardless, both B cells and MDCs showed that they had upregulated the activation markers CD80, CD70 and lymph node homing marker CCR7 after MAB273 or CP-870,893 exposure (Figures 1C and 1D). B cell proliferation, as assessed by dilution of CellTrace violet dye, also showed similar activation with MAB273 or CP-870,893 stimulation (Figure 1E). Although induction of phenotypic differentiation and cell proliferation was clear with both MAB273 and CP-870,893, we did not find detectable cytokines such as IL-12 p40, IL-6, IFN-γ and TNF in the cell culture supernatants after stimulation (data not shown).

**Figure 1.**
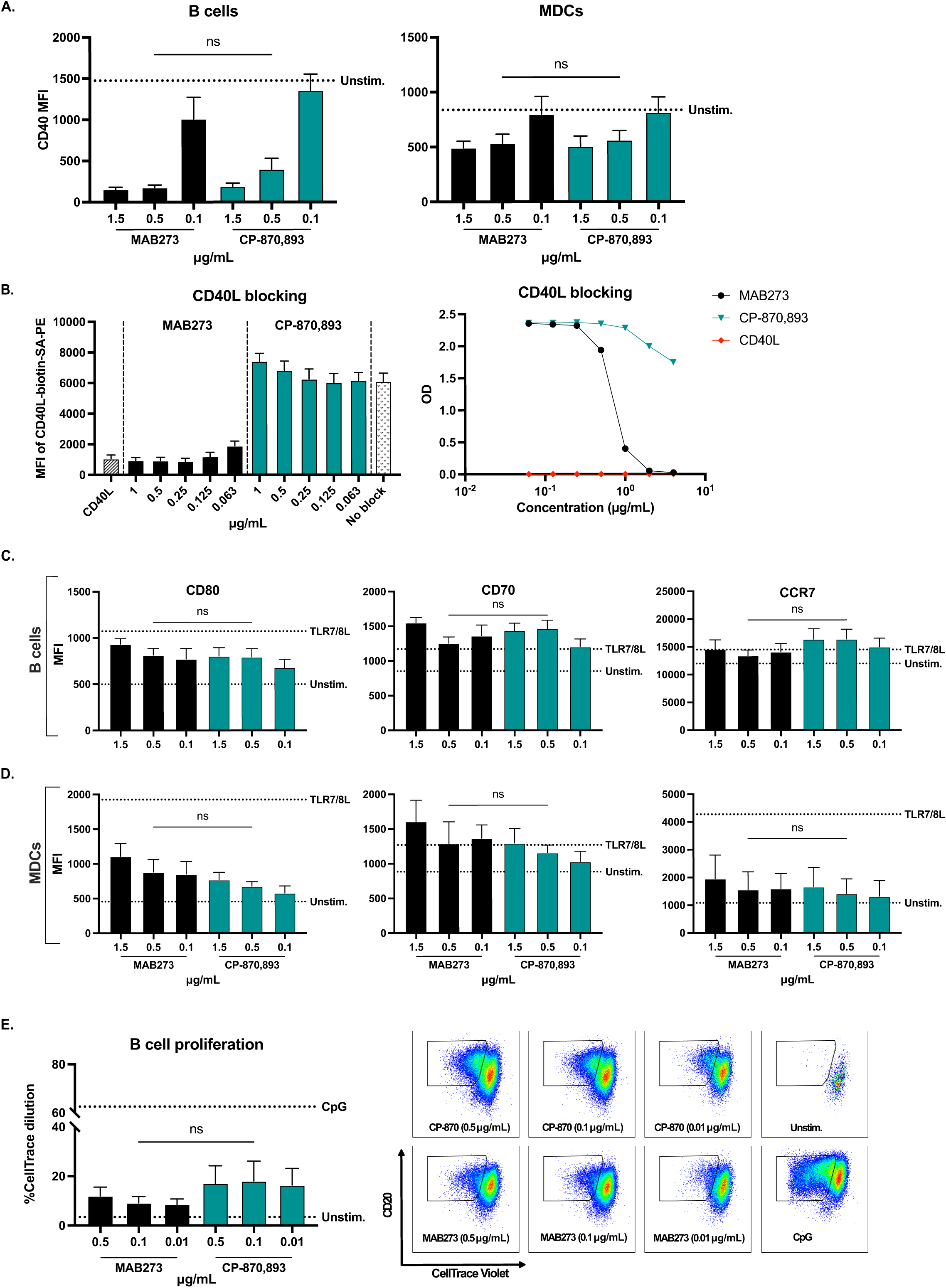
MAB273 binds CD40 and activates human B cells and MDCs with a similar potency as CP-870,893 *in vitro*. n=3, mean ± SEM. **(A).** Surface expression levels of CD40 were evaluated by flow cytometry after exposure with anti-CD40 Abs (1.5, 0.5, 0.1 µg/mL) for 2 hours at 4°C. **(B).** Flow cytometry (left) and ELISA (right) results showing the signal of competitive CD40L-biotin-streptavidin conjugate after culture with anti-CD40 Abs (1, 0.5, 0.25, 0.125, 0.063 µg/mL) or CD40L (2 µg/mL) for 20 minutes at 4°C (for flow cytometry) or 2 hours at room temperature (for ELISA). **(C-D).** Human PBMCs were stimulated with anti-CD40 Abs (1.5, 0.5, 0.1 µg/mL) or TLR7/8L (5 µg/mL) for 24 hours at 37°C. Cell activation markers CD80, CD70 and LN homing marker CCR7 on B cells **(C)** and MDCs **(D)** were evaluated by flow cytometry. **(E).** Human PBMCs were stimulated with anti-CD40 Abs (0.5, 0.1, 0.01 µg/mL) or CpG (1 µg/mL) for 5 days. B cell proliferation was indicated by percentage of CellTrace Violet negative B cells. Representative flow cytometry plots are shown. ns = not statistically significant. See also Figure S1.

### CD40 binding and activation capacities remain after removing the Fc region of MAB273

In order to evaluate if MAB273 is FcγR-independent and the role of avidity, we generated F(ab’)2 fragments by pepsin digestion to cleave off the Fc region but maintain the hinge region as well as Fab fragments by papain digestion to remove both the Fc region and hinge region (Figure 2A). We confirmed that the Fab and F(ab’)2 fragments of MAB273 still bound to CD40 but were not detected by an anti-Fc antibody (Figure 2B). Human B cells and MDCs exposed to MAB273 or the F(ab’)2 fragment showed similar ability to block the CD40 staining antibody in a dose dependent manner, while the Fab fragment showed weaker CD40 blocking capacity (Figure 2C). In addition, MAB273 and the F(ab’)2 exposure resulted in similar upregulation of CD80, CD70 and CCR7 on B cells (Figure 2D) and MDCs (Figure 2E), while the Fab fragment retained the activation capacity on MDCs (Figure 2E) but was weaker for B cells (Figure 2D) suggesting that B cells may require higher avidity for activation (Figure S1A). In addition, B cell proliferation induced by MAB273 or F(ab’)2 was similar but Fab showed weaker induction (Figure 2F). This demonstrates that the F(ab’)2 fragment of MAB273 has retained immunostimulatory capacities *in vitro* after removing the Fc region, but the Fab fragment was less potent for B stimulation. Nevertheless, we conclude that MAB273-induced activation is not dependent on FcγR crosslinking.

**Figure 2.**
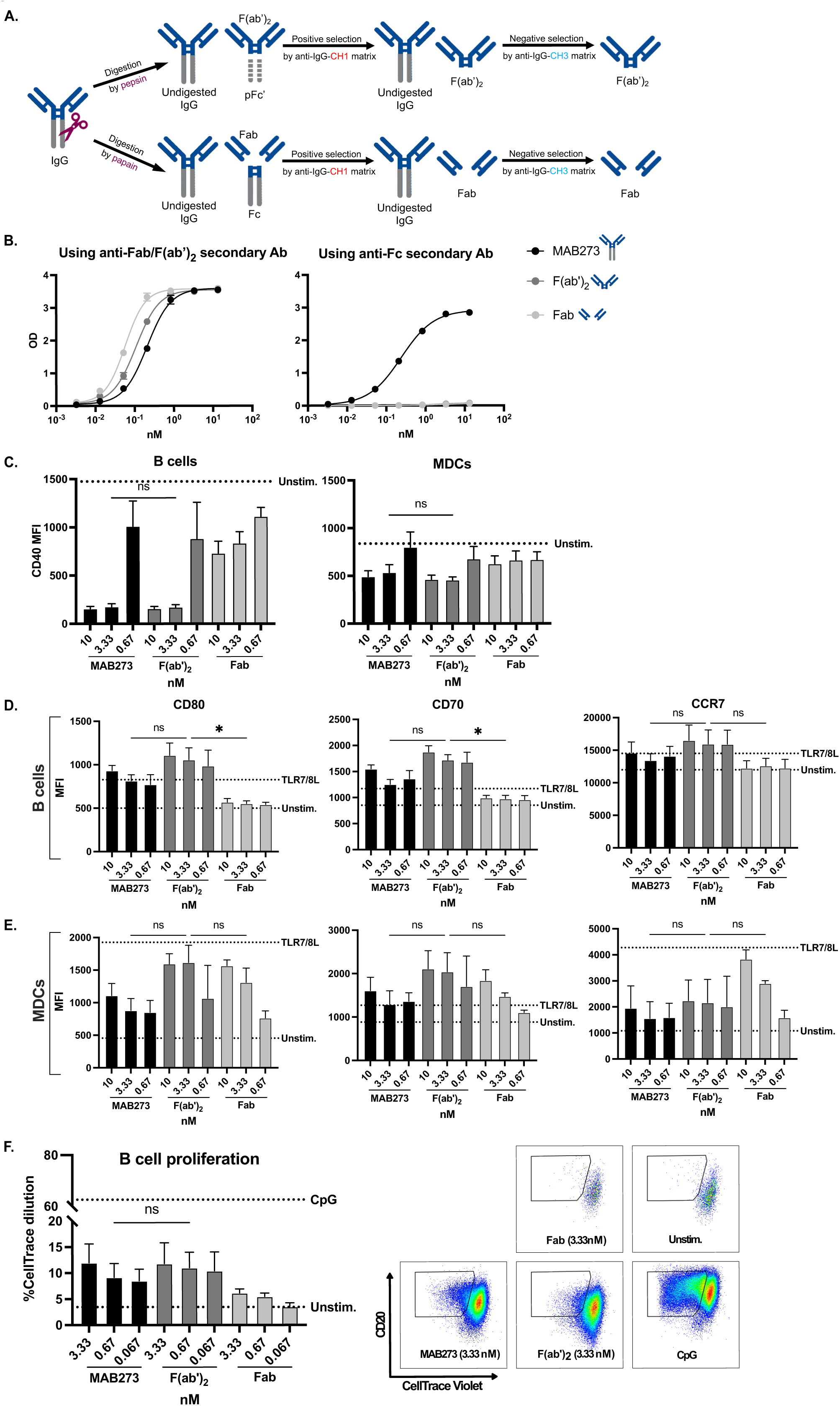
CD40 binding and activation remain the same after removing the Fc region of MAB273. n=3, mean ± SEM. **(A).** Cartoon showing the process of generating Fab and F(ab’)2 fragments of MAB273. **(B).** ELISA results showing the CD40 binding capacity while using anti-Fab/F(ab’)2 (left) or anti-Fc (right) secondary Abs. **(C).** Surface expression levels of CD40 were evaluated by flow cytometry after exposure with complete MAB273 or its Fab/F(ab’)2 fragments (10, 3.33, 0.67 nM) for 2 hours at 4°C. **(D-E).** Human PBMCs were stimulated with complete MAB273, its Fab/F(ab’)2 fragments (10, 3.33, 0.67 nM) or TLR7/8L (5 µg/mL, as positive control) for 24 hours at 37°C. Cell activation markers CD80, CD70 and LN homing marker CCR7 on B cells **(D)** and MDCs **(E)** were evaluated by flow cytometry. **(F).** Human PBMCs were stimulated with complete MAB273, its Fab/F(ab’)2 fragments (3.33, 0.67, 0.067 nM) or CpG (1 µg/mL, as positive control) for 5 days. B cell proliferation was indicated by percentage of CellTrace Violet negative B cells. Representative flow cytometry plots are shown. *p < 0.05, ns = not statistically significant. See also Figure S1.

### MAB273 binds CD40 and activates rhesus macaque PBMCs in vitro

With the further aim of utilizing a physiological *in vivo* animal model, we next tested the ability of MAB273 to bind and stimulate rhesus macaque PBMCs *in vitro* by repeating a large subset of the above *in vitro* experiments. The immune cells expressed CD40 as expected where B cells and MDCs had the highest expression and T cells and neutrophils low expression (Figure S1B). Phenotypic differentiation after stimulation with MAB273, CP-870,893 or the isotype control antibody (IgG1-LALA) were analyzed. Again, the signal of the CD40 staining antibody was blocked when the cells had been exposed to MAB273 or CP-870,893 but not to the isotype control (Figures 3A and 3B). In addition, MAB273 upregulated CD80 on B cells (Figure 3A) and MDCs (Figure 3B). Less upregulation was found by CP-870,893 and no upregulation was found by the isotype control antibody. B cell proliferation was also induced by MAB273 exposure but not by the isotype control antibody (Figure 3C). MAB273 induced low but detectable levels of IL-12 p40, IL-6 and TNF secretion (Figure 3D). IFN-γ was not detected (data not shown). This shows that MAB273 can bind to rhesus macaque CD40 and activate immune cells.

**Figure 3.**
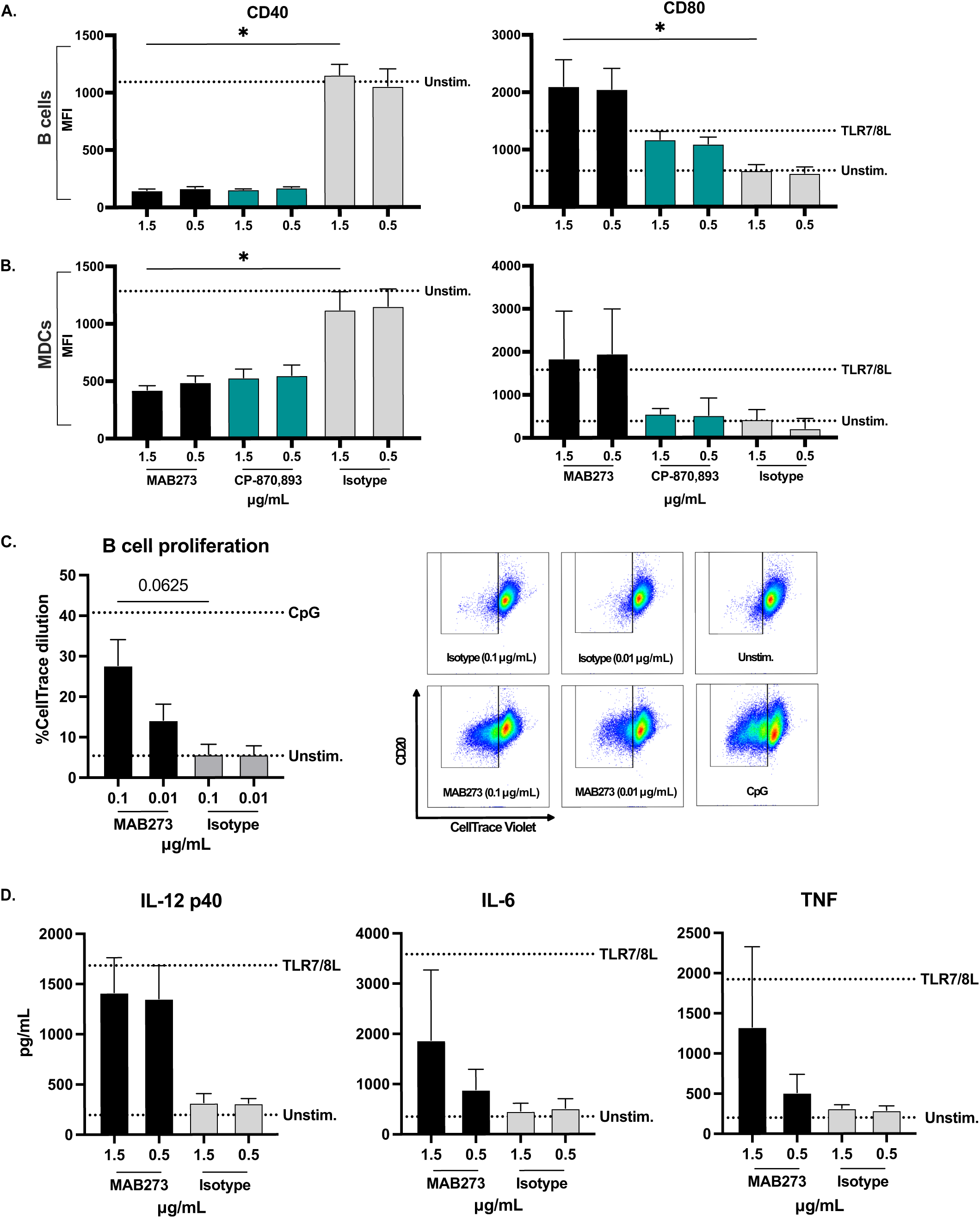
MAB273 shows potent CD40 binding and activation capacities in rhesus macaque PBMCs *in vitro*. Rhesus PBMCs were stimulated with anti-CD40 Abs, isotype control Ab (1.5, 0.5 µg/mL) for 2 hours at 4°C (for CD40) or additional TLR7/8L (5 µg/mL, as positive control) for 24 hours at 37°C (for CD80). Surface expression levels of CD40 and cell activation marker CD80 on B cells **(A)** and MDCs **(B)** were evaluated by flow cytometry. n=6, mean ± SEM. **(C).** Rhesus PBMCs were stimulated with MAB273, isotype control Ab (0.1, 0.01 µg/mL) or CpG (1 µg/mL, as positive control) for 5 days. B cell proliferation was indicated by percentage of CellTrace Violet negative B cells. Representative flow cytometry plots are shown. n=5, mean ± SEM. **(D).** Level of cytokines (IL-12 p40, IL-6 and TNF) were measured by ELISA, supernatants used were taken from **(A)** and **(B)**. n=3, mean ± SEM. *p < 0.05. See also Figure S1.

### MAB273 induces innate immune activity in vivo in rhesus macaques

Six rhesus macaques were thereafter divided into three groups to receive MAB273 administration at different doses and routes (Figure 4A). Standard clinical chemistry analyses, including a series of liver and kidney function and complete blood count (CBC) measurements were performed in addition to immunological analyses. The first group that received the highest dose of 1 mg/kg by intravenous (i.v.) administration showed that several clinical chemistry parameters including alkaline phosphatase (ALP), alanine transaminase (ALT), gamma-glutamyl transferase (GGT), bile acid (BA), total bilirubin (TBIL) and blood urea nitrogen (BUN) were elevated above the normal reference range (Figure S2A). This group also showed side effects characterized by loss of appetite and reduced activity behavior for up to 5 days. In contrast, the lower dose of 0.1 mg/kg given i.v. did not induce any detectable side effects and most of the clinical chemistry parameters remained within the healthy range (Figure S2A). The low dose was thereafter tested with subcutaneous (s.c.) administration which neither induced side effects. As visualized by CBC, there were fluctuations of cell numbers following administration of MAB273 found with both doses and routes. A rapid decline in platelets was observed already at 0.5-4 hours accompanied by a rapid increase in white blood cell counts, especially granulocytes, was found after MAB273 administration in all groups (Figure S2B). Frequencies of specific cell subsets identified by flow cytometry and normalized to the CBC data confirmed a rapid increase in neutrophils while there was a transient decline in both B cells and MDCs. The cell fluctuations were dose-dependent and with a notably more dramatic effect in the 1 mg/kg i.v. group (Figure S2C). The transient fluctuation of immune cells after MAB273 administration may stem from redistribution of activated cells leaving the circulation to migrate to tissues followed by a replenishment of cells from the bone marrow as has been proposed earlier[11, 22]. Body weight remained stable during the entire study period in all groups (Figure S2D).

**Figure 4.**
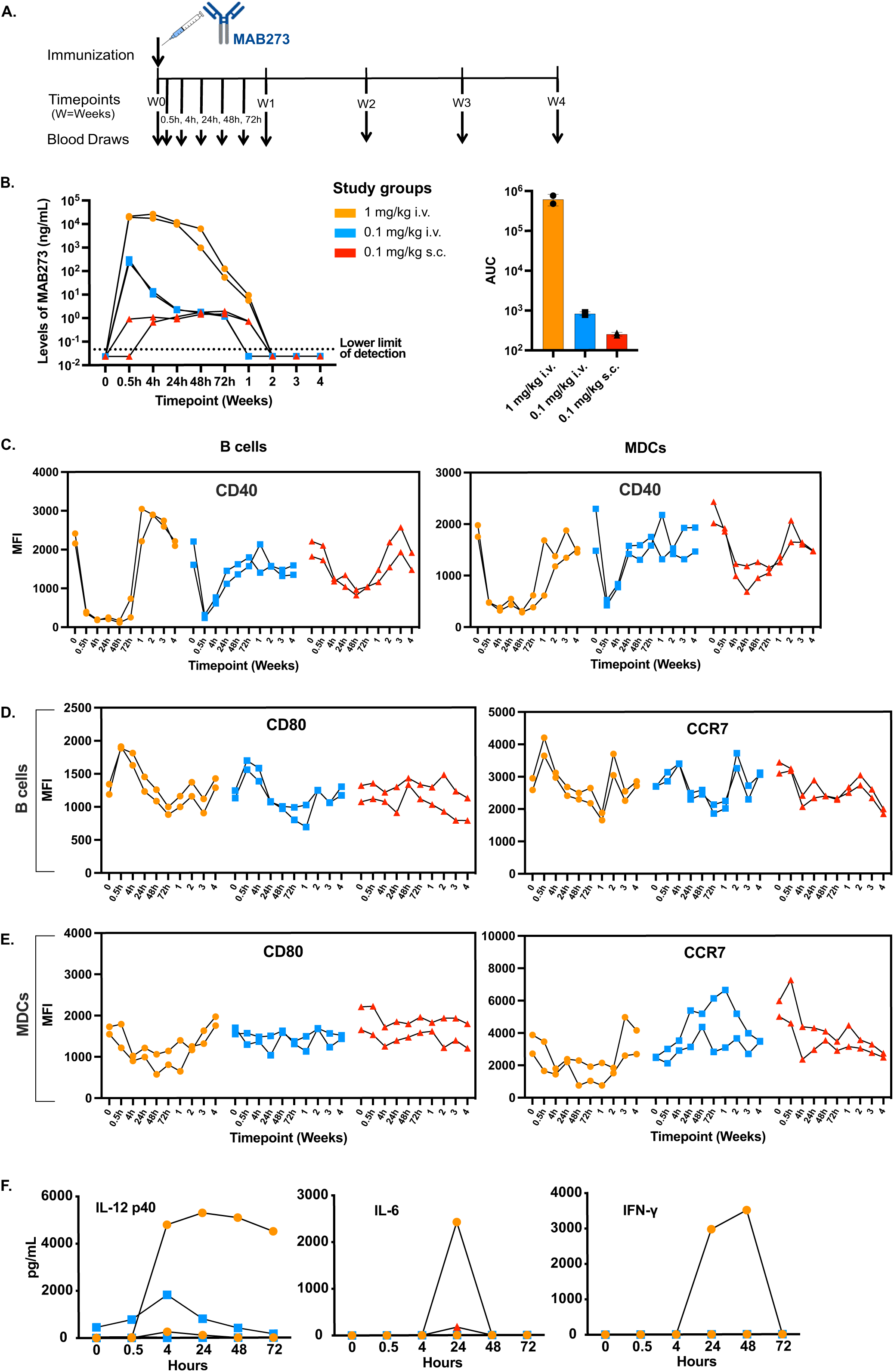
*In vivo* innate immune activity in rhesus macaques. **(A).** Outline for toxicity and safety study in rhesus macaques administered 1 mg/kg i.v., 0.1 mg/kg i.v., or 0.1 mg/kg s.c. (n=2 per group). **(B).** Levels of MAB273 in plasma over time (left), AUC (area under curve, right) is calculated after normalizing (left) to linear axes. n=2 per group. **(C-E).** Surface expression levels of CD40 **(C)**, cell activation markers CD80 and lymph node homing marker CCR7 on B cells **(D)** and MDCs **(E)** were evaluated by flow cytometry over time. n=2 per group. **(F)** Systemic levels of pro-inflammatory cytokines (IFN-γ, IL-6, IL-12 p40) in plasma over time. n=2 per group. See also Figure S2.

Analysis of the pharmacokinetics (PK) of MAB273 in plasma showed that the levels were readily detectable after 0.5 hour of administration in both of the i.v. groups (Figure 4B). In the high dose group, the levels of MAB273 peaked around 0.5-4 hours and then declined gradually until it was undetectable after 2 weeks. In the low dose group, the highest level was detected at 0.5 hour, then continually decreased and was undetectable after 1 week. In the s.c. group, MAB273 was detectable at 0.5-4 hours but at 2-4 log lower levels compared to the i.v. groups. However, the level of MAB273 in the s.c. group was sustained for a week and undetectable at 2 weeks. This suggests that s.c. administration results in a depot effect and slower release of MAB273 into the circulation.

Binding of MAB273 to CD40 *in vivo* was evaluated by quantifying the loss of detection signal from the staining CD40 antibody as performed in the *in vitro* experiments. Rapidly (0.5 hour) after administration of MAB273, detection of CD40 was blocked on B cells and MDCs (Figure 4C). Lack of CD40 signal was sustained for 72 hours in the high dose i.v. group while this was found for a shorter period for the low dose i.v. group. The s.c. group also showed reduced signal for CD40, but this was noticed later (at 4 hours) and sustained for 2 weeks in line with the observed pharmacokinetics of MAB273 in plasma. As mentioned above, the return of detectable CD40 expression may be due to replenishment of new cells into the circulation as well as the half-life of MAB273. Accompanied with MAB273 binding to CD40 on immune cells, a rapid increase in CD80 and CCR7 expression was observed especially in i.v. groups (Figure 4D). The expression gradually returned to baseline levels or even below which may be explained by that newly recruited cells exhibit a more immature phenotype. The upregulation of CD80 and CCR7 on MDCs was less noticeable than on B cells (Figure 4E). Secretion of IL-12 p40, IL-6 and IFN-γ was detected in one of the animals receiving the high dose while most animals did not show detectable levels (Figure 4F). TNF was not detected (data not shown). In conclusion, MAB273 induces strong innate immune activation with regards to cell recruitment and activation while being well-tolerated at the dose of 0.1 mg/kg given either i.v. or s.c. in rhesus macaques. Since s.c. administration demonstrated clear immune stimulation while potentially offering a depot effect of MAB273 for slower release and better tolerability, this route may be more attractive for clinical development and hence this was used in our subsequent studies.

### MAB273 targets and activates immune cells at the site of injection and draining lymph nodes

To understand the biodistribution of MAB273 in different tissues after administration, the antibody was labeled with AlexaFluor 680 fluorochrome to enable tracking *in vivo*. The fluorescent signal and unaltered CD40 binding and activation capacities of the labeled MAB273 were validated *in vitro* before administered *in vivo* (Figures S3A-S3C). Three animals were immunized s.c. and biopsies were collected after 24 (n=1) or 48 hours (n=2) from the sites of injection, lymph nodes (LNs) and other selected tissues (Figure 5A). MAB273-AlexaFluor 680 was predominantly detected at the site of injection (skin of the left thigh) and the LNs specifically draining this site (left inguinal LNs, left common iliac LNs and paraaortic LNs). Monocytes, B cells, neutrophils, MDCs, PDCs and macrophages showed detectable MAB273 binding while T cells had no signal (Figure 5B). MAB273 signal was not detected at the saline control injection site in the skin of the opposite thigh (right) and the LNs draining this site (right inguinal LNs and right common iliac LNs). No or very weak signal was detected in draining peripheral tissues such as the liver, spleen, bone marrow, BAL and PBMCs (Figure 5B and S3D). There were more CD45+ immune cells targeted with MAB273 at the injection site and the primary LNs compared to the secondary draining LNs more distant from the injection site (Figure 5C). In the skin, the most abundant cell subsets targeted with MAB273 were macrophages, neutrophils and monocytes likely due to CD40 expression and phagocytic ability[30–32]. In the draining LNs, B cells were predominantly targeted with MAB273 likely due to that they represent a major CD40 expressing population[33–35] (Figure 5D). In line with the detectable signal of the MAB273, there was considerable infiltration of immune cells to the injection site in the skin compared to the saline-injection sites. This consisted of mainly infiltrating B cells, MDCs, monocytes, neutrophils and T cells (Figures 3E-3G). Again, reduced CD40 staining signal was observed at the injection site, draining LNs and PBMCs indicative of MAB273 binding to CD40 expressing cells as expected (Figures 5E and 5F). We therefore concluded that MAB273 has restricted biodistribution to the site of injection and specific draining LNs and targets multiple cell subsets but to the highest degree B cells, DCs and macrophages.

**Figure 5.**
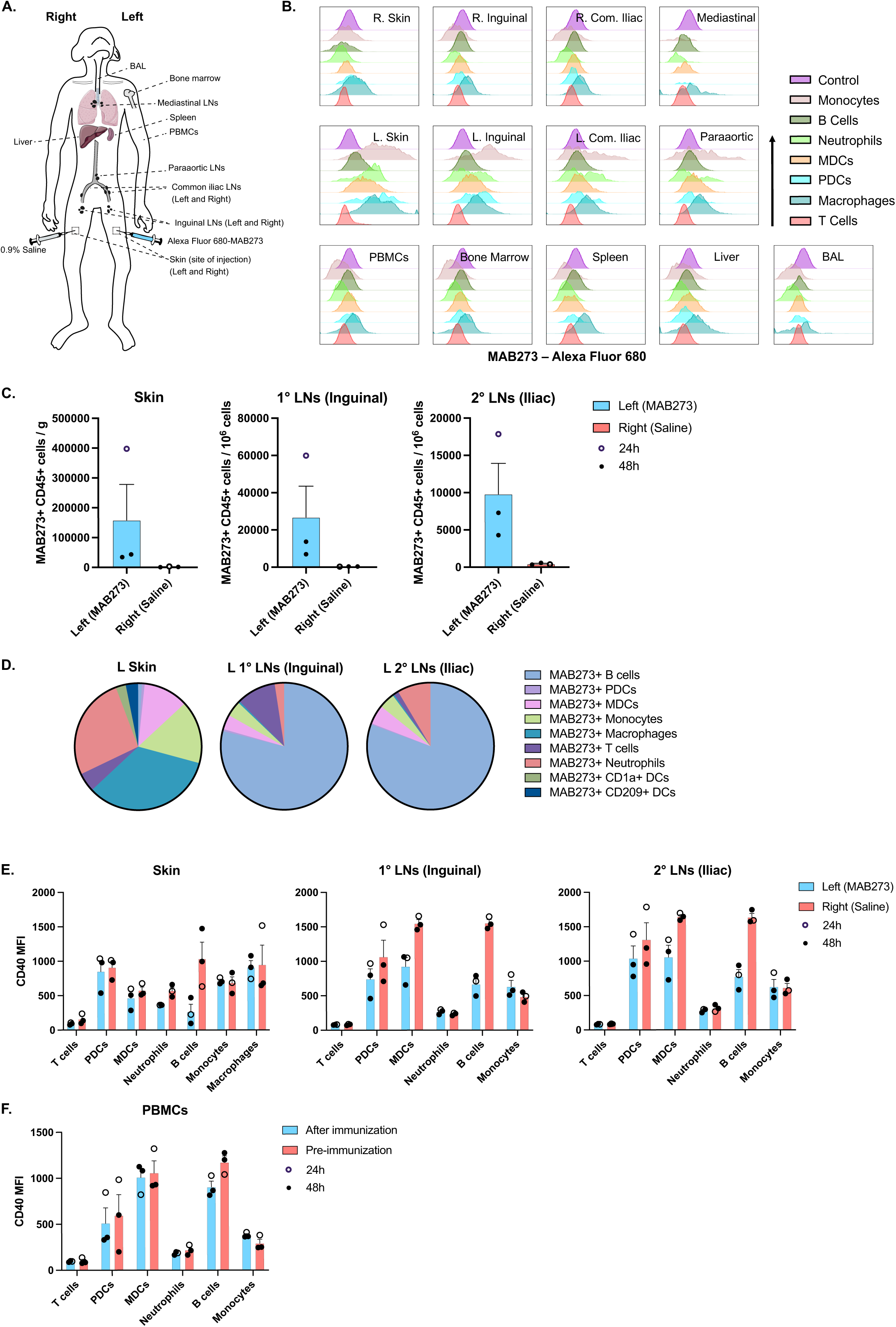
*In vivo* biodistribution of MAB273. **(A).** Rhesus macaques (n=3) were administered 0.1mg/kg s.c. of Alexa Fluor 680-MAB273 in the skin above the left quad and 0.9% saline solution s.c. in the skin above the right quad. The cartoon shows the sites of immunization and sampling performed after 24 hours (n=1) or 48 hours (n=2). **(B).** Histograms show Alexa Fluor 680-MAB273 signal on different cell populations in different tissues of one representative animal. Control = peripheral blood B cells from the same animal before immunization with labeled MAB273. **(C).** MAB273+ CD45+ cells normalized by counting beads at site of injection or draining lymph nodes. n=3, mean ± SEM. **(D).** Pie charts show proportion of different CD45+ immune cells targeted with MAB273 at the injection site, the primary and secondary draining LNs. **(E-F).** Expression of CD40 at site of injection and draining lymph nodes **(E)** as well as PBMCs **(F)**. Compiled data were evaluated by flow cytometry. Geometric mean fluorescence intensity (MFI) is shown. n=3, mean ± SEM. See also Figure S3.

### Strong induction of genes associated with innate immune stimulation in MAB273 targeted tissues

To further understand the immune activation profile induced by MAB273 administration *in vivo,* we performed RNA sequencing analyses on the draining LNs and skin from the site of injection as well as the blood (Figure 6A). This revealed a significant number of differentially expressed genes (DEGs) in the MAB273 targeted skin and LNs compared to the donor-matched saline control sites (Figures 6B and 6C). In addition, blood taken before MAB273 administration compared to 24-48 hours after showed significant gene modulation (Figure 6D).

**Figure 6.**
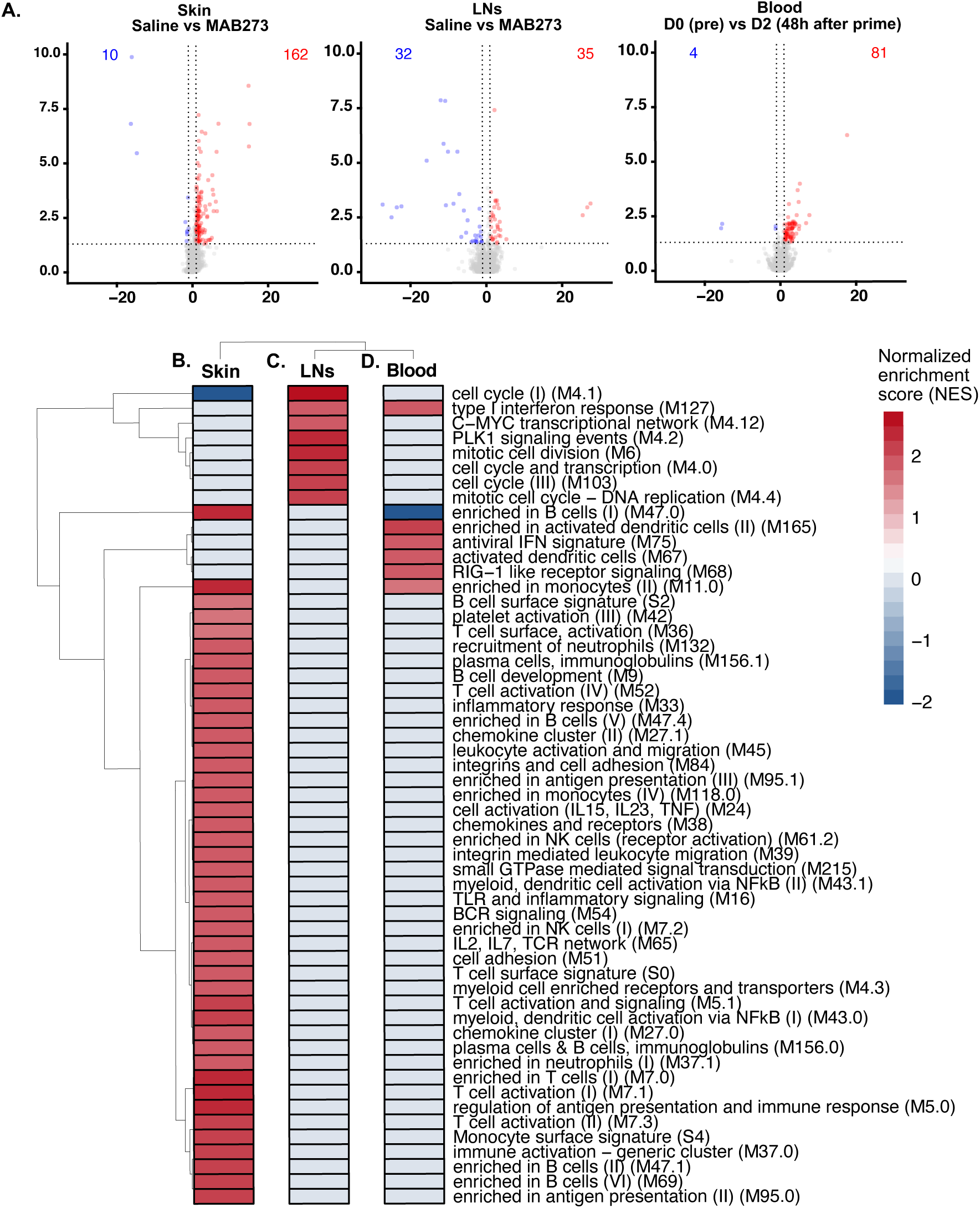
RNA-seq data analysis in different tissues. RNA-sequencing was performed on samples from rhesus macaques administered with AF680-MAB273 (see Figure 5). n=3. **(A).** Volcano plots of differentially expressed genes in skin, inguinal lymph nodes and blood. Up-regulated genes are in red and down-regulated genes are in blue. The calculation of differentially expressed genes was based on a control reference for each tissue: D0 pre-immunization (blood) or saline site of injection (lymph nodes and skin). Dotted grey lines indicate fold change > 1 and adjusted p-values < 0.05. **(B-D).** Gene seat enrichment analysis (GSEA) of blood transcription modules significantly enriched in skin **(B)**, lymph nodes **(C)** and blood **(D)** compared to their respective control samples, color gradient is based on normalized gene set enrichment scores. All statistical comparisons were adjusted by the Benjamini-Hochberg procedure, adjusted p-values < 0.05 were considered significant. “D0” is the day before prime immunization, “D2” is 48h after prime. See also Figure S3.

Gene set enrichment analysis using the blood transcription modules (BTMs) described previously[29] demonstrated that distinctly different gene modules were changed at the different anatomical sites. The skin had the highest activation and transcriptional changes after MAB273 injection. The upregulated genes in skin included sets of genes associated with specific cell surface markers (*CD19*, *CD2, IL21R)* and chemokines such as the *CXCR5* gene. All the significantly enriched gene modules were upregulated, except the cell cycle modules. The results indicated activation and recruitment of T cells, B cells, NK cells, monocytes, and DCs to the site of injection (Figure 6B). MAB273-draining LNs also showed that there were genes upregulated compared to the saline-draining LNs. These genes were fewer and were distinct from those observed in the skin. The upregulated genes in the LNs were mainly linked to antigen presentation (*IRAG2*) and interferon (*IRF6*), and a few downregulated genes were linked to RNA processing (*U2*, *U3*, *U4*, *RNaseP*). The enrichment analysis indicated an upregulation of modules related to cell proliferation (mitotic cell division and cell cycle modules) (Figure 6C). Furthermore, in the blood, the genes differentially expressed 24-48 hours after MAB273 administration were mainly associated with interferon signatures, such as *ISG15*, *RSAD2*, and *SKIV2L* as well as genes associated with monocytes and DC activation (Figure 6D). MAB273 therefore induces significant innate immune activation characterized by monocyte and DC activation in the blood, recruitment of immune cells to the site of injection while cell proliferation and antigen presentation processes were more dominant in the draining LNs.

### MAB273 exhibits adjuvant effects for induction of antigen-specific CD4 and CD8 T cells

Finally, we evaluated the effect of MAB273 to act as an adjuvant both for therapeutic vaccination where low degree of immunity already exists and also to enhance primary immune responses as in prophylactic vaccination. Three animals therefore first received seven well characterized HIV-1 envelope glycoprotein (Env) peptides[24] as model antigen alone two times to establish low levels of immunity before receiving boost immunizations with MAB273 co-administered s.c. to mimic a therapeutic vaccination (Figure 7A). In a separate group, three animals received MAB273 together with the Env peptides in a prime-boost schedule of four immunizations to mimic prophylactic vaccination. The final immunization was performed with an additional recombinant trimer Env protein (Figure 7A).

**Figure 7.**
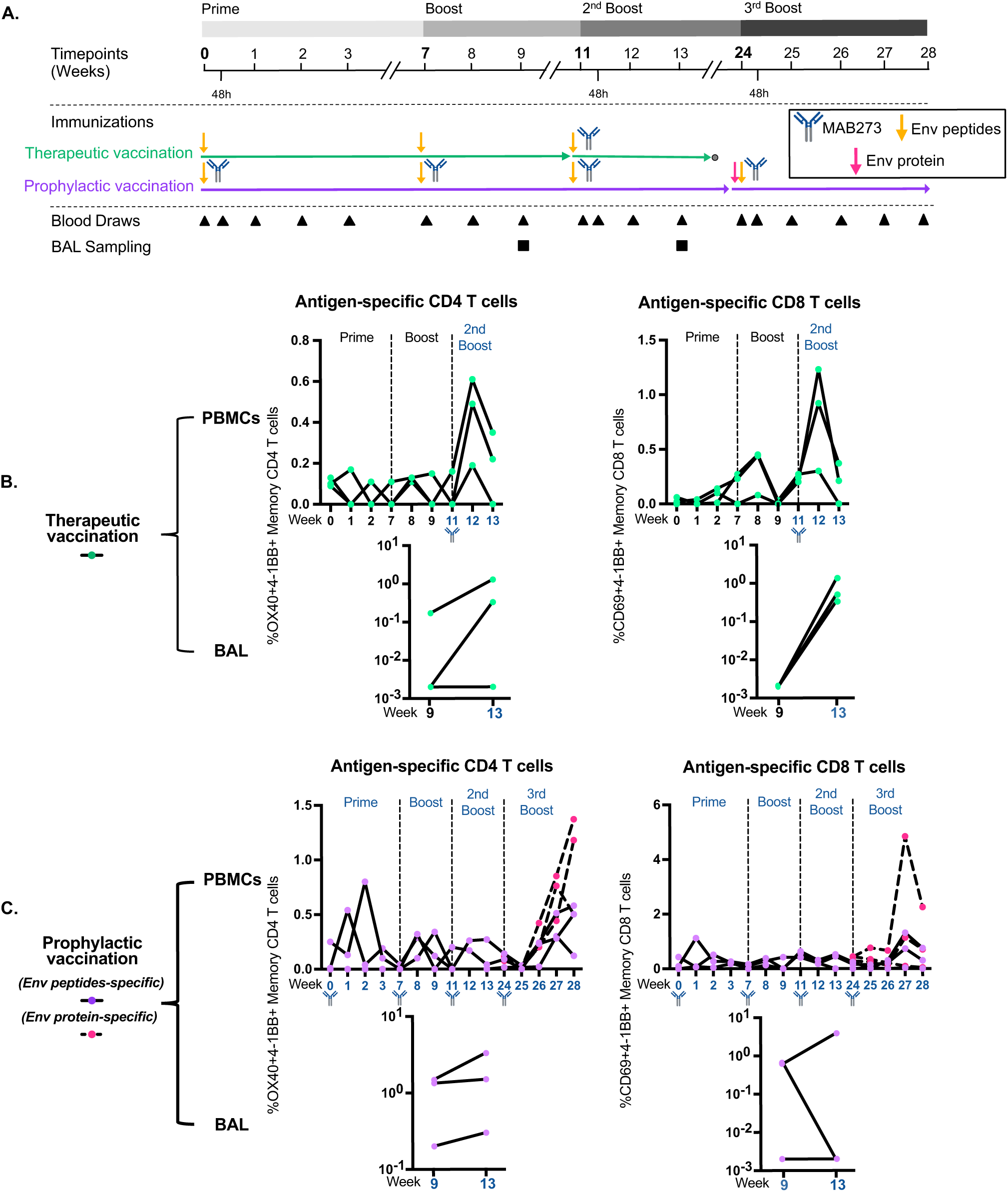
MAB273 can potentiate induction of antigen-specific CD4 T cells and CD8 T cells in PBMCs and BAL. **(A).** In the therapeutic vaccination group, rhesus macaques (n=3) were administered 0.1mg/kg s.c. of Env peptides for the first two immunizations then 1 mg/kg Env peptides plus 0.1 mg/kg MAB273 s.c. for the last immunization; in the prophylactic vaccination group, rhesus macaques (n=3) were co-injected with 0.1 mg/kg Env peptides plus 0.1 mg/kg MAB273 s.c. for the first two immunizations then 1 mg/kg Env peptides plus 0.1 mg/kg MAB273 s.c. for the third immunization, followed 1 mg/kg Env peptides, 0.1 mg/kg MAB273 plus 100 μg Env protein for the last immunization. **(B).** Antigen-specific CD4 T cells and CD8 T cells in PBMCs and BAL in therapeutic vaccination group. n=3. **(C).** Antigen-specific CD4 T cells and CD8 T cells in PBMCs and BAL in prophylactic vaccination group. n=3. See also Figure S4.

Low frequencies of Env-specific T cell responses were induced by Env peptide immunization alone (Figure 7B). The responses were enhanced in two out of three animals when they received a boost with Env peptides and MAB273. This effect was evident for both systemic Env-specific CD4 and CD8 T cells in blood (Figure 7B) as well as in bronchoalveolar lavage (BAL) (Figure 7B). Two out of the three animals immunized with Env peptides in combination with MAB273 already at prime immunization induced higher levels of Env-specific CD4 and CD8 T cell responses compared to the animals receiving Env peptides only (Figures 7B and 7C). Although the subsequent boost immunizations re-activated T cell responses, they did not reach the peak levels found after the prime immunization (Figure 7C). This was observed both in blood and BAL and may be a consequence of the low dose of Env peptides (0.1 mg/kg) and the induction of antibodies against the humanized MAB273 in rhesus macaques (Figures S4A and S4B).

The activation profile of MAB273 based on the RNA sequencing and blood transcriptome analysis comparing the activation at pre-immunization compared to the second boost showed that the differences were negligible indicating that recurrent MAB273 administration may result in lower innate immune activation (Figure S4C). Nevertheless, reactivation of memory T cell responses to peak levels occurred after the fourth immunization of MAB273 when using trimer Env protein in combination with Env peptides to provide more antigen (Figure 7C). No detectable IgG to Env peptides was found (Figure S4D) but IgG to Env protein was detected (Figure S4E). Taken together, our study demonstrates that MAB273 is a potent agonistic anti-CD40 antibody with rapid binding and activation to B cells and MDCs *in vitro* and *in vivo* in rhesus macaques and can help enhance antigen-specific T cell responses.

## Discussion

This study provides evidence that MAB273 binds to CD40 and activates human and rhesus macaque B cells and MDCs *in vitro* and in rhesus macaques *in vivo.* In particular, MAB273 activates CD40 signaling to upregulate T cell costimulatory receptors CD80 and CD70 and LN homing receptor CCR7 on APCs which aids in driving T cell responses as shown by induction of both systemic (blood) and tissue (BAL) immunity. This is in line with what has been reported with other potent anti-CD40 agonistic antibodies[11, 22, 26, 36–39]. However, MAB273 binds to the CD40L binding site of CD40 and exerts its biological activity independent of FcγR crosslinking which is a unique combined feature.

Structure analysis has demonstrated that a symmetric complex between trimeric CD40L and dimers of CD40 is formed when they interact[40, 41]. CD40 signaling requires large clusters of these complexes[42, 43]. However, CD40L has also been reported to interact with several integrins independent of CD40-CD40L interaction[44, 45]. In fact, multiple CD40L binding sites for integrins may form even larger (anti-CD40 antibody)-CD40-CD40L-intergrin complexes that induce additional biological activities than the canonical CD40 activation which could enhance unwanted side effects[46–50]. Based on this, we screened and developed a series of antibodies, including MAB273, which can effectively replace CD40L and potentially avoid interference by additional CD40-CD40L-intergrin complex formation[23]. Our results confirmed that MAB273 binds to the CD40L binding site. Earlier studies have shown that APX005M, a mAb competing with the CD40L binding site, is highly agonistic and may activate CD40 similarly to endogenous CD40L[22, 51].

CD40 signaling induced by agonistic anti-CD40 antibodies has been shown to depend on cells expressing Fc-receptors[18]. In this regard, engineering the Fc region of agonistic CD40 antibodies to promote the FcγR crosslinking can enhance their potency[18–22], but increased crosslinking also augments adverse events[20, 21]. However, agonistic anti-CD40 antibodies of the IgG2 isotype show Fc-independent activity[38, 52]. This is likely provided by conformational regulation and flexibility of disulfide bonds in the hinge region[52, 53] which facilitates CD40 clustering[42, 43] in contrast to the IgG1 isotype which has an unfixed hinge region and highly flexible Fab arms. However, there are also opposite results showing that the activity of IgG2 isotype antibody is not Fc-independent[20]. Taken together, at least in terms of the IgG1 isotype, functional Fc region and FcγR crosslinking are considered necessary for strong CD40 agonists. However, our results with MAB273 demonstrate, both *in vitro* and *in vivo*, that it is possible for agonistic CD40 antibodies to be of the IgG1 isotype and function independently of FcγR crosslinking[16]. We have earlier found that the Fc-silenced MAB273 induced more potent immune activity than several variants of CP-870,893, including the strongest Fc-enhanced crosslinking antibody CP-870,893 IgG1-V11[23], which supports the notion that epitope binding site is critical. Moreover, in this study we observed that only bivalent F(ab’)2, and not monovalent Fab, showed similar agonistic activity as complete MAB273. This suggests that clustering of CD40 by bivalent Fab arms is a critical component of activation and may explain why IgG2 agonists retain activity without Fc engagement[39, 42, 52]. Still, various published agonistic CD40 antibodies show different or even completely opposite functions in terms of CD40L-binding site specificity and FcγR crosslinking[11, 16, 22, 37, 38].

We tested MAB273 in rhesus macaques in order to mimic the human immune system as closely as possible. This was partly driven by that agonistic CD40 antibodies have distinct characteristics in mice and humans[16] and human FcγRs are different from mouse FcγRs[54]. Previous studies have mostly used human CD40 transgenic (hCD40Tg) mice[37, 55], hCD40Tg FcγRIIb-/- mice[52], hCD40Tg FcγR-/- mice[43] or hCD40Tg/mFcgr2b-/- /hFcgr2b+/- mice[42]. However, it has been proposed that only humanized CD40/FcγR mice can provide the correct *in vivo* environment for evaluating CD40 antibodies[20]. On the other hand, despite that macaque FcγRIIb binds poorly to human antibodies[54], non-human primates (NHPs) are the preferred preclinical model[22, 26, 27, 38] due to their immunophenotypic similarity and CD40 homology to humans. To this end, a preclinical study of an Fc-unmodified CD40 agonist, CDX-1140, showed that the agonistic activity is Fc-independent and well-tolerated in NHPs[38]. However, the Fc region of CDX-1140 is still functional and thus NHPs may not accurately predict activity, toxicity, or Fc-independence in humans. Since MAB273 has double LALA (L234A and L235A)-mutations to completely eliminate Fc-FcγR binding, the NHP model should largely reflect the activity in humans in this regard. This is also supported by that CD40-binding and activation in human and rhesus cells showed similar results *in vitro*. Previous studies demonstrated that LALA critically reduces the binding of the Fc region to all known FcγRs and the complement component 1q (C1q). As a result, antibody-dependent cellular cytotoxicity (ADCC) and complement-dependent cytotoxicity (CDC) are not induced[56–58]. LALA has no effect on serum clearance[59], nor on PK[60].

Our dose escalation results showed that 1 mg/kg i.v. gave side effects by transient elevation of liver transaminases and behavioral changes like reduced appetite and physical activity, while 0.1 mg/kg did not but still induced robust immune activation. The peak of liver transaminases appeared on day 7 and normalized by day 21 which is delayed compared to results reported for CP-870,893, which appeared between day 2-8[11]. We noted that MAB273 induced transient liver abnormality similar to reported by other agonistic CD40 antibodies[11, 21, 26] which may be caused by apoptosis of CD40 expressing hepatocytes and CD40-mediated hyperactivation[11, 61]. However, hepatic toxicity can often be controlled by dose and route of administration as indicated by our results showing that the dose of 0.1 mg/kg did not cause changes in liver function. By tracking fluorescently labeled MA273 after s.c. administration, we also observed almost no fluorescent signal in the liver. We have earlier shown that i.v. administration of another CD40 antibody resulted on distribution in the liver[26, 27]. The transient increase of blood urea nitrogen (BUN) in 1 mg/kg i.v. group could be explained by the difficulty of renal excretion induced by macromolecular drugs (such as antibodies) and consequential renal inflammatory responses[62]. Transient hematologic changes were also observed after administration and aligned well with the pharmacokinetics of MAB273 in all groups in our study. In particular, the number of B cells and MDCs decreased in blood. Also, a rapid decline in platelets at 0.5 hour was found, likely caused by activation of CD40-expressing platelets and their contribution to inflammation and aggregation[44, 63]. Our *in vitro* and RNA sequencing results also suggested that MAB273 can induce B cell proliferation similarly to other studies[26, 37, 38]. The downregulation of the B cell enrichment module in the blood and concomitant upregulation in the skin followed the change in cell numbers and suggest an extravasation and replenishment of immune cells induced by CD40 activation, as has been proposed earlier[11, 22, 38, 42]. The rapid increase in numbers of granulocytes, especially neutrophils, at 0.5 hour to 4 hours likely contributed to most of the observed enrichment in inflammation signatures. The 1 mg/kg i.v. group showed overall higher magnitude and longer duration of inflammation but 0.1 mg/kg given s.c. induced larger fluctuations in cell numbers in blood than 0.1 mg/kg i.v.. This may be caused by that s.c. administration also stimulated cells locally in the skin which resulted in more redistribution of neutrophils[64].

Anti-drug antibodies (ADA), such as the anti-MAB273 IgG observed in our study, are antibodies raised to the administered human antibodies and have often not been controlled for in earlier CD40 antibody studies. Induction of ADA has frequently been reported with a variety of other human antibodies administered to macaques[65–67]. ADA may result in reduced potency of the CD40 antibodies during subsequent administrations and affect the accuracy of conclusions drawn from sequential dosing, even when using humanized CD40/FcγR mice as the animal model. The ADA we observed after each immunization may have interfered with the efficiency of the boost immunizations. Nevertheless, boosting of antigen-specific T cell responses could be detected after each immunization and especially when a higher antigen dose including Env protein was given. Additional studies using more animals and an optimal dose of antigen, perhaps as well as using a rhesus version of MAB273, are needed to assess the enhancement of T cell responses by the adjuvant effect of MAB273.

In summary, our study shows the safety, cell targeting and immunostimulatory properties of this novel agonistic anti-CD40 antibody of IgG1 isotype that is CD40L binding site specific and works independently of FcγR crosslinking. These are distinct features from previously reported agonistic anti-CD40 antibodies and may therefore offer new avenues for the adjuvant targeting of the CD40:CD40L pathway.

## Acknowledgements

We thank Richard Wyatt, Shridhar Bale and Richard Wilson (Scripps Research, California, United States) for kindly providing the 426c NFL trimer of Env protein.

## Author contributions

Conceptualization – X.Y., S.O., J.F., D.P., U.P., S.F., K.L.; Formal Analysis – X.Y., S.O., R.A., K.Y., K.L.; Funding acquisition – J.F., D.P., U.P., S.F., K.L.; Investigation – X.Y., S.O., R.A., K.Y., K.Le., F.H., A.C.; Methodology – X.Y., S.O., R.A., K.Le., F.H., A.C., K.Y., M.B., F.N., J.F., D.P., U.P., S.F., K.L.; Resources – F.N., J.F., D.P., U.P., S.F., K.L.; Supervision – K.L.; Visualization – X.Y., R.A., K.Y.; Writing – original draft – X.Y., K.L.; Writing – review & editing – all authors.

## Funding

This work was supported by Icano MAB GmbH and grants from the Swedish Research Council (Vetenskapsrådet; 2019-01036 and 2020-05829 to K.L.). This research was also supported by a grant from the China Scholarship Council (X.Y.) and intramural faculty salary grants from Karolinska Institutet (S.O., K.Le., F.H.).

## Data availability

The datasets generated during and/or analysed during the current study are available from the corresponding author on reasonable request. The manuscript has data included as electronic supplementary material.

## Declaration of interests statement

J.F., D.P., U.P., S.F. are employees of Icano MAB GmbH. All other authors declare no competing interests.

---

**Supplementary Figure 1. Baseline expression of CD40. Related to Figure 1-3. (A).** Baseline expression of CD40 on different human immune cells. Mean fluorescence intensity (MFI) of CD40 is shown (n=7). **(B).** Baseline expression of CD40 on different rhesus macaque immune cells. MFI of CD40 is shown (n=13). **(C).** Gating strategy for innate phenotyping.

**Supplementary Figure 2. Safety monitoring and change of cell frequencies. Related to Figure 4. (A).** Safety monitoring by clinical chemistry tests. **(B).** Safety monitoring by complete blood counts. **(C).** Cell frequencies normalized by CBC over time. **(D).** Weight and body temperature change over time.

**Supplementary Figure 3. Validation of Alexa Fluor 680-conjugated MAB273 and gating strategy. Related to Figure 5**. Alexa Fluor 680-conjugated MAB273 labeling test **(A)**, binding test **(B)** and activation test **(C)** before tracking immunization. **(D).** Gating strategy for tracking Alexa Fluor 680-conjugated MAB273.

**Supplementary Figure 4. MAB273 signal *in vivo* and immune cell infiltration. Related to Figure 5 and 6. (A).** MAB273+ CD45+ cells normalized by counting beads in peripheral tissues and blood. n=3, mean ± SEM. **(B).** CD45+ immune cells normalized by weight or counting beads at site of injection and draining lymph nodes. **(C).** Graphs show cell subsets from **(B)**. n=3, mean ± SEM. **(D).** Pie charts show proportion of different CD45+ immune cells at the injection site, the primary and secondary draining LNs. **(E).** Flow plots of one representative animal show MAB273+ B cells and MAB273+ MDCs at site of injection and draining lymph nodes as well as PBMCs.

**Supplementary Figure 5. Antibody responses after MAB273 administration. Related to Figure 7. (A).** Levels of MAB273 in plasma over time in immunogenicity study. LLOQ=lower limit of quantification, LLOD=lower limit of detection. n=3. **(B).** Rhesus anti-human MAB273 IgG titers over time. **(C).** Volcano plots of differentially expressed genes in blood, “D0” is the day before prime immunization, “D79” is 48h after the second boost. **(D).** Anti-Env peptides IgG titers over time. **(E).** Anti-Env protein IgG titers over time. Rhesus Plasma from animals who received six immunizations with Env protein immunization was used as positive control.

**Supplementary Table 1.**
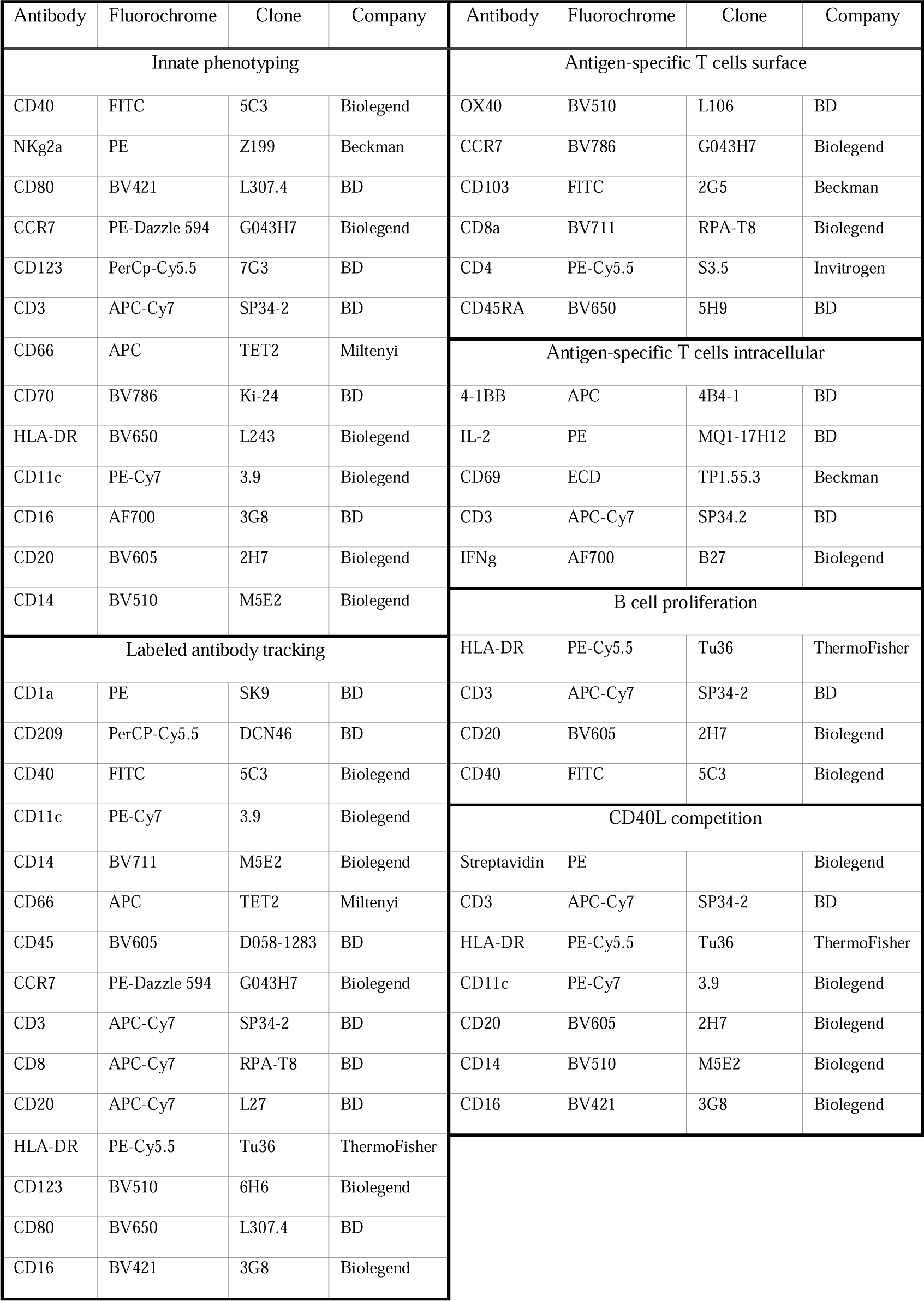
Fluorescent staining antibodies used in flow cytometry.

## References

1. Zhang H, Dai Z, Wu W, Wang Z, Zhang N, Zhang L, et al (2021) Regulatory mechanisms of immune checkpoints PD-L1 and CTLA-4 in cancer. J Exp Clin Cancer Res 40(1):184. https://doi.org/10.1186/s13046-021-01987-7

2. Rowshanravan B, Halliday N, Sansom DM (2018) CTLA-4: a moving target in immunotherapy. Blood 131(1):58–67. https://doi.org/10.1182/blood-2017-06-741033

3. Boutros C, Tarhini A, Routier E, Lambotte O, Ladurie FL, Carbonnel F, et al (2016) Safety profiles of anti-CTLA-4 and anti-PD-1 antibodies alone and in combination. Nat Rev Clin Oncol 13(8):473–86. https://doi.org/10.1038/nrclinonc.2016.58

4. Vonderheide RH (2018) The Immune Revolution: A Case for Priming, Not Checkpoint. Cancer Cell 33(4):563–9. https://doi.org/10.1016/j.ccell.2018.03.008

5. Elgueta R, Benson MJ, de Vries VC, Wasiuk A, Guo Y, Noelle RJ (2009) Molecular mechanism and function of CD40/CD40L engagement in the immune system. Immunol Rev 229(1):152–72. https://doi.org/10.1111/j.1600-065X.2009.00782.x

6. van Kooten C, Banchereau J (2000) CD40-CD40 ligand. J Leukoc Biol 67(1):2–17. https://doi.org/10.1002/jlb.67.1.2

7. French RR, Chan HT, Tutt AL, Glennie MJ (1999) CD40 antibody evokes a cytotoxic T-cell response that eradicates lymphoma and bypasses T-cell help. Nat Med 5(5):548–53. https://doi.org/10.1038/8426

8. Bennett SR, Carbone FR, Karamalis F, Flavell RA, Miller JF, Heath WR (1998) Help for cytotoxic-T-cell responses is mediated by CD40 signalling. Nature 393(6684):478–80. https://doi.org/10.1038/30996

9. Diehl L, den Boer AT, Schoenberger SP, van der Voort EI, Schumacher TN, Melief CJ, et al (1999) CD40 activation in vivo overcomes peptide-induced peripheral cytotoxic T-lymphocyte tolerance and augments anti-tumor vaccine efficacy. Nat Med 5(7):774–9. https://doi.org/10.1038/10495

10. Vonderheide RH (2020) CD40 Agonist Antibodies in Cancer Immunotherapy. Annu Rev Med 71:47–58. https://doi.org/10.1146/annurev-med-062518-045435

11. Vonderheide RH, Flaherty KT, Khalil M, Stumacher MS, Bajor DL, Hutnick NA, et al (2007) Clinical activity and immune modulation in cancer patients treated with CP-870,893, a novel CD40 agonist monoclonal antibody. J Clin Oncol 25(7):876–83. https://doi.org/10.1200/jco.2006.08.3311

12. Beatty GL, Torigian DA, Chiorean EG, Saboury B, Brothers A, Alavi A, et al (2013) A phase I study of an agonist CD40 monoclonal antibody (CP-870,893) in combination with gemcitabine in patients with advanced pancreatic ductal adenocarcinoma. Clin Cancer Res 19(22):6286–95. https://doi.org/10.1158/1078-0432.Ccr-13-1320

13. Vonderheide RH, Burg JM, Mick R, Trosko JA, Li D, Shaik MN, et al (2013) Phase I study of the CD40 agonist antibody CP-870,893 combined with carboplatin and paclitaxel in patients with advanced solid tumors. Oncoimmunology 2(1):e23033. https://doi.org/10.4161/onci.23033

14. Bajor DL, Xu X, Torigian DA, Mick R, Garcia LR, Richman LP, et al (2014) Immune activation and a 9-year ongoing complete remission following CD40 antibody therapy and metastasectomy in a patient with metastatic melanoma. Cancer Immunol Res 2(11):1051–8. https://doi.org/10.1158/2326-6066.Cir-14-0154

15. Rüter J, Antonia SJ, Burris HA, Huhn RD, Vonderheide RH (2010) Immune modulation with weekly dosing of an agonist CD40 antibody in a phase I study of patients with advanced solid tumors. Cancer Biol Ther 10(10):983–93. https://doi.org/10.4161/cbt.10.10.13251

16. Richman LP, Vonderheide RH (2014) Role of crosslinking for agonistic CD40 monoclonal antibodies as immune therapy of cancer. Cancer Immunol Res 2(1):19–26. https://doi.org/10.1158/2326-6066.Cir-13-0152

17. White AL, Chan HT, French RR, Beers SA, Cragg MS, Johnson PW, et al (2013) FcγRΙΙB controls the potency of agonistic anti-TNFR mAbs. Cancer Immunol Immunother 62(5):941–8. https://doi.org/10.1007/s00262-013-1398-6

18. White AL, Chan HT, Roghanian A, French RR, Mockridge CI, Tutt AL, et al (2011) Interaction with FcγRIIB is critical for the agonistic activity of anti-CD40 monoclonal antibody. J Immunol 187(4):1754–63. https://doi.org/10.4049/jimmunol.1101135

19. Li F, Ravetch JV (2011) Inhibitory Fcγ receptor engagement drives adjuvant and anti-tumor activities of agonistic CD40 antibodies. Science 333(6045):1030–4. https://doi.org/10.1126/science.1206954

20. Dahan R, Barnhart BC, Li F, Yamniuk AP, Korman AJ, Ravetch JV (2016) Therapeutic Activity of Agonistic, Human Anti-CD40 Monoclonal Antibodies Requires Selective FcγR Engagement. Cancer Cell 29(6):820–31. https://doi.org/10.1016/j.ccell.2016.05.001

21. Knorr DA, Dahan R, Ravetch JV (2018) Toxicity of an Fc-engineered anti-CD40 antibody is abrogated by intratumoral injection and results in durable antitumor immunity. Proc Natl Acad Sci U S A 115(43):11048–53. https://doi.org/10.1073/pnas.1810566115

22. Filbert EL, Björck PK, Srivastava MK, Bahjat FR, Yang X (2021) APX005M, a CD40 agonist antibody with unique epitope specificity and Fc receptor binding profile for optimal therapeutic application. Cancer Immunol Immunother 70(7):1853–65. https://doi.org/10.1007/s00262-020-02814-2

23. Reitinger C, Beckmann K, Carle A, Bluemle E, Jurkschat N, Paulmann C, et al (2023) Fcgamma-receptor-independent controlled activation of CD40 canonical signaling by novel therpeutic antibodies for cancer therapy. bioRxiv:2023.01.16.521736. https://doi.org/10.1101/2023.01.16.521736

24. Nehete PN, Nehete BP, Hill L, Manuri PR, Baladandayuthapani V, Feng L, et al (2008) Selective induction of cell-mediated immunity and protection of rhesus macaques from chronic SHIV(KU2) infection by prophylactic vaccination with a conserved HIV-1 envelope peptide-cocktail. Virology 370(1):130–41. https://doi.org/10.1016/j.virol.2007.08.022

25. Lenart K, Hellgren F, Ols S, Yan X, Cagigi A, Cerveira RA, et al (2022) A third dose of the unmodified COVID-19 mRNA vaccine CVnCoV enhances quality and quantity of immune responses. Mol Ther Methods Clin Dev 27:309–23. https://doi.org/10.1016/j.omtm.2022.10.001

26. Thompson EA, Liang F, Lindgren G, Sandgren KJ, Quinn KM, Darrah PA, et al (2015) Human Anti-CD40 Antibody and Poly IC:LC Adjuvant Combination Induces Potent T Cell Responses in the Lung of Nonhuman Primates. J Immunol 195(3):1015–24. https://doi.org/10.4049/jimmunol.1500078

27. Thompson EA, Darrah PA, Foulds KE, Hoffer E, Caffrey-Carr A, Norenstedt S, et al (2019) Monocytes Acquire the Ability to Prime Tissue-Resident T Cells via IL-10-Mediated TGF-β Release. Cell Rep 28(5):1127–35.e4. https://doi.org/10.1016/j.celrep.2019.06.087

28. Ols S, Yang L, Thompson EA, Pushparaj P, Tran K, Liang F, et al (2020) Route of Vaccine Administration Alters Antigen Trafficking but Not Innate or Adaptive Immunity. Cell Rep 30(12):3964–71.e7. https://doi.org/10.1016/j.celrep.2020.02.111

29. Li S, Rouphael N, Duraisingham S, Romero-Steiner S, Presnell S, Davis C, et al (2014) Molecular signatures of antibody responses derived from a systems biology study of five human vaccines. Nat Immunol 15(2):195–204. https://doi.org/10.1038/ni.2789

30. Buhtoiarov IN, Lum H, Berke G, Paulnock DM, Sondel PM, Rakhmilevich AL (2005) CD40 ligation activates murine macrophages via an IFN-gamma-dependent mechanism resulting in tumor cell destruction in vitro. J Immunol 174(10):6013–22. https://doi.org/10.4049/jimmunol.174.10.6013

31. Beatty GL, Chiorean EG, Fishman MP, Saboury B, Teitelbaum UR, Sun W, et al (2011) CD40 agonists alter tumor stroma and show efficacy against pancreatic carcinoma in mice and humans. Science 331(6024):1612–6. https://doi.org/10.1126/science.1198443

32. Salomon R, Rotem H, Katzenelenbogen Y, Weiner A, Cohen Saban N, Feferman T, et al (2022) Bispecific antibodies increase the therapeutic window of CD40 agonists through selective dendritic cell targeting. Nat Cancer 3(3):287–302. https://doi.org/10.1038/s43018-022-00329-6

33. De Silva NS, Klein U (2015) Dynamics of B cells in germinal centres. Nat Rev Immunol 15(3):137–48. https://doi.org/10.1038/nri3804

34. Mesin L, Ersching J, Victora GD (2016) Germinal Center B Cell Dynamics. Immunity 45(3):471–82. https://doi.org/10.1016/j.immuni.2016.09.001

35. Young C, Brink R (2021) The unique biology of germinal center B cells. Immunity 54(8):1652–64. https://doi.org/10.1016/j.immuni.2021.07.015

36. Carpenter EL, Mick R, Rüter J, Vonderheide RH (2009) Activation of human B cells by the agonist CD40 antibody CP-870,893 and augmentation with simultaneous toll-like receptor 9 stimulation. J Transl Med 7:93. https://doi.org/10.1186/1479-5876-7-93

37. Mangsbo SM, Broos S, Fletcher E, Veitonmäki N, Furebring C, Dahlén E, et al (2015) The human agonistic CD40 antibody ADC-1013 eradicates bladder tumors and generates T-cell-dependent tumor immunity. Clin Cancer Res 21(5):1115–26. https://doi.org/10.1158/1078-0432.Ccr-14-0913

38. Vitale LA, Thomas LJ, He LZ, O’Neill T, Widger J, Crocker A, et al (2019) Development of CDX-1140, an agonist CD40 antibody for cancer immunotherapy. Cancer Immunol Immunother 68(2):233–45. https://doi.org/10.1007/s00262-018-2267-0

39. Yu X, Chan HTC, Orr CM, Dadas O, Booth SG, Dahal LN, et al (2018) Complex Interplay between Epitope Specificity and Isotype Dictates the Biological Activity of Anti-human CD40 Antibodies. Cancer Cell 33(4):664–75.e4. https://doi.org/10.1016/j.ccell.2018.02.009

40. Naismith JH, Devine TQ, Brandhuber BJ, Sprang SR (1995) Crystallographic evidence for dimerization of unliganded tumor necrosis factor receptor. J Biol Chem 270(22):13303–7. https://doi.org/10.1074/jbc.270.22.13303

41. Vanamee É S, Faustman DL (2018) Structural principles of tumor necrosis factor superfamily signaling. Sci Signal 11(511). https://doi.org/10.1126/scisignal.aao4910

42. Yu X, Chan HTC, Fisher H, Penfold CA, Kim J, Inzhelevskaya T, et al (2020) Isotype Switching Converts Anti-CD40 Antagonism to Agonism to Elicit Potent Antitumor Activity. Cancer Cell 37(6):850–66.e7. https://doi.org/10.1016/j.ccell.2020.04.013

43. Yu X, James S, Felce JH, Kellermayer B, Johnston DA, Chan HTC, et al (2021) TNF receptor agonists induce distinct receptor clusters to mediate differential agonistic activity. Commun Biol 4(1):772. https://doi.org/10.1038/s42003-021-02309-5

44. Aloui C, Prigent A, Sut C, Tariket S, Hamzeh-Cognasse H, Pozzetto B, et al (2014) The signaling role of CD40 ligand in platelet biology and in platelet component transfusion. Int J Mol Sci 15(12):22342–64. https://doi.org/10.3390/ijms151222342

45. El Fakhry Y, Alturaihi H, Yacoub D, Liu L, Guo W, Leveillé C, et al (2012) Functional interaction of CD154 protein with α5β1 integrin is totally independent from its binding to αIIbβ3 integrin and CD40 molecules. J Biol Chem 287(22):18055–66. https://doi.org/10.1074/jbc.M111.333989

46. McWhirter SM, Pullen SS, Holton JM, Crute JJ, Kehry MR, Alber T (1999) Crystallographic analysis of CD40 recognition and signaling by human TRAF2. Proc Natl Acad Sci U S A 96(15):8408–13. https://doi.org/10.1073/pnas.96.15.8408

47. An HJ, Kim YJ, Song DH, Park BS, Kim HM, Lee JD, et al (2011) Crystallographic and mutational analysis of the CD40-CD154 complex and its implications for receptor activation. J Biol Chem 286(13):11226–35. https://doi.org/10.1074/jbc.M110.208215

48. Smulski CR, Beyrath J, Decossas M, Chekkat N, Wolff P, Estieu-Gionnet K, et al (2013) Cysteine-rich domain 1 of CD40 mediates receptor self-assembly. J Biol Chem 288(15):10914–22. https://doi.org/10.1074/jbc.M112.427583

49. Léveillé C, Bouillon M, Guo W, Bolduc J, Sharif-Askari E, El-Fakhry Y, et al (2007) CD40 ligand binds to alpha5beta1 integrin and triggers cell signaling. J Biol Chem 282(8):5143–51. https://doi.org/10.1074/jbc.M608342200

50. Tong AW, Stone MJ (2003) Prospects for CD40-directed experimental therapy of human cancer. Cancer Gene Ther 10(1):1–13. https://doi.org/10.1038/sj.cgt.7700527

51. O’Hara MH, O’Reilly EM, Varadhachary G, Wolff RA, Wainberg ZA, Ko AH, et al (2021) CD40 agonistic monoclonal antibody APX005M (sotigalimab) and chemotherapy, with or without nivolumab, for the treatment of metastatic pancreatic adenocarcinoma: an open-label, multicentre, phase 1b study. Lancet Oncol 22(1):118–31. https://doi.org/10.1016/s1470-2045(20)30532-5

52. White AL, Chan HT, French RR, Willoughby J, Mockridge CI, Roghanian A, et al (2015) Conformation of the human immunoglobulin G2 hinge imparts superagonistic properties to immunostimulatory anticancer antibodies. Cancer Cell 27(1):138–48. https://doi.org/10.1016/j.ccell.2014.11.001

53. Orr CM, Fisher H, Yu X, Chan CH, Gao Y, Duriez PJ, et al (2022) Hinge disulfides in human IgG2 CD40 antibodies modulate receptor signaling by regulation of conformation and flexibility. Sci Immunol 7(73):eabm3723. https://doi.org/10.1126/sciimmunol.abm3723

54. Bournazos S, DiLillo DJ, Ravetch JV (2015) The role of Fc-FcγR interactions in IgG-mediated microbial neutralization. J Exp Med 212(9):1361–9. https://doi.org/10.1084/jem.20151267

55. Ceglia V, Zurawski S, Montes M, Kroll M, Bouteau A, Wang Z, et al (2021) Anti-CD40 Antibody Fused to CD40 Ligand Is a Superagonist Platform for Adjuvant Intrinsic DC-Targeting Vaccines. Front Immunol 12:786144. https://doi.org/10.3389/fimmu.2021.786144

56. Lund J, Pound JD, Jones PT, Duncan AR, Bentley T, Goodall M, et al (1992) Multiple binding sites on the CH2 domain of IgG for mouse Fc gamma R11. Mol Immunol 29(1):53–9. https://doi.org/10.1016/0161-5890(92)90156-r

57. Tamm A, Schmidt RE (1997) IgG binding sites on human Fc gamma receptors. Int Rev Immunol 16(1-2):57–85. https://doi.org/10.3109/08830189709045703

58. Bournazos S, Gupta A, Ravetch JV (2020) The role of IgG Fc receptors in antibody-dependent enhancement. Nat Rev Immunol 20(10):633–43. https://doi.org/10.1038/s41577-020-00410-0

59. Wines BD, Powell MS, Parren PW, Barnes N, Hogarth PM (2000) The IgG Fc contains distinct Fc receptor (FcR) binding sites: the leukocyte receptors Fc gamma RI and Fc gamma RIIa bind to a region in the Fc distinct from that recognized by neonatal FcR and protein A. J Immunol 164(10):5313–8. https://doi.org/10.4049/jimmunol.164.10.5313

60. Leabman MK, Meng YG, Kelley RF, DeForge LE, Cowan KJ, Iyer S (2013) Effects of altered FcγR binding on antibody pharmacokinetics in cynomolgus monkeys. MAbs 5(6):896–903. https://doi.org/10.4161/mabs.26436

61. Afford SC, Randhawa S, Eliopoulos AG, Hubscher SG, Young LS, Adams DH (1999) CD40 activation induces apoptosis in cultured human hepatocytes via induction of cell surface fas ligand expression and amplifies fas-mediated hepatocyte death during allograft rejection. J Exp Med 189(2):441–6. https://doi.org/10.1084/jem.189.2.441

62. Meibohm B, Zhou H (2012) Characterizing the impact of renal impairment on the clinical pharmacology of biologics. J Clin Pharmacol 52(1 Suppl):54s–62s. https://doi.org/10.1177/0091270011413894

63. Inwald DP, McDowall A, Peters MJ, Callard RE, Klein NJ (2003) CD40 is constitutively expressed on platelets and provides a novel mechanism for platelet activation. Circ Res 92(9):1041–8. https://doi.org/10.1161/01.Res.0000070111.98158.6c

64. Liew PX, Kubes P (2019) The Neutrophil’s Role During Health and Disease. Physiol Rev 99(2):1223–48. https://doi.org/10.1152/physrev.00012.2018

65. Gardner MR, Fetzer I, Kattenhorn LM, Davis-Gardner ME, Zhou AS, Alfant B, et al (2019) Anti-drug Antibody Responses Impair Prophylaxis Mediated by AAV-Delivered HIV-1 Broadly Neutralizing Antibodies. Mol Ther 27(3):650–60. https://doi.org/10.1016/j.ymthe.2019.01.004

66. Lee WS, Reynaldi A, Amarasena T, Davenport MP, Parsons MS, Kent SJ (2021) Anti-Drug Antibodies in Pigtailed Macaques Receiving HIV Broadly Neutralising Antibody PGT121. Front Immunol 12:749891. https://doi.org/10.3389/fimmu.2021.749891

67. Kim YJ, Koh EM, Song CH, Byun MS, Choi YR, Jeon EJ, et al (2021) Preclinical immunogenicity testing using anti-drug antibody analysis of GX-G3, Fc-fused recombinant human granulocyte colony-stimulating factor, in rat and monkey models. Sci Rep 11(1):12004. https://doi.org/10.1038/s41598-021-91360-7

