## Supplementary Figures for "Cell targeting and immunostimulatory properties of a novel Fcγ-receptor independent agonistic anti-CD40 antibody in rhesus macaques"

Supplementary Figure 1

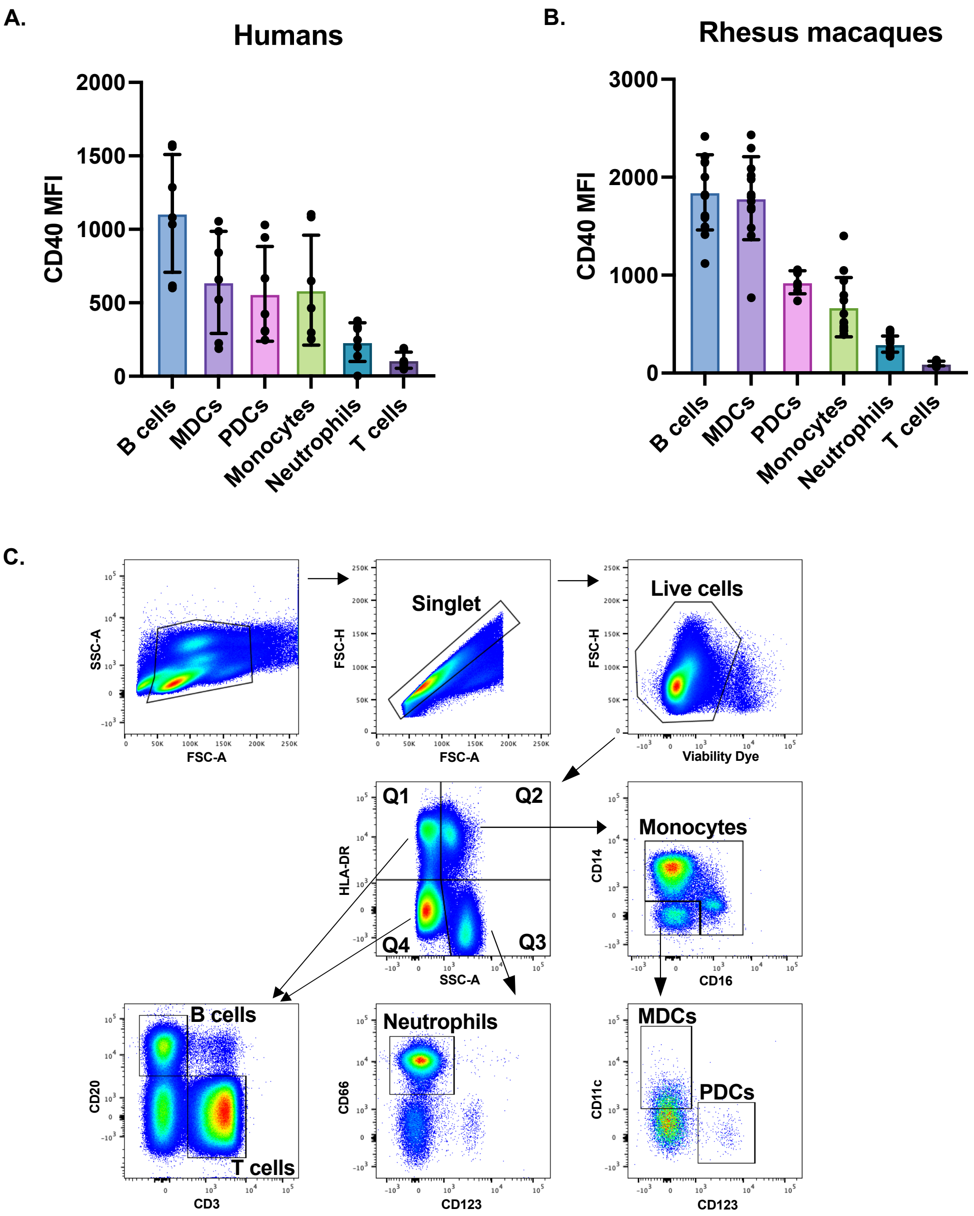

**Supplementary Figure 2**

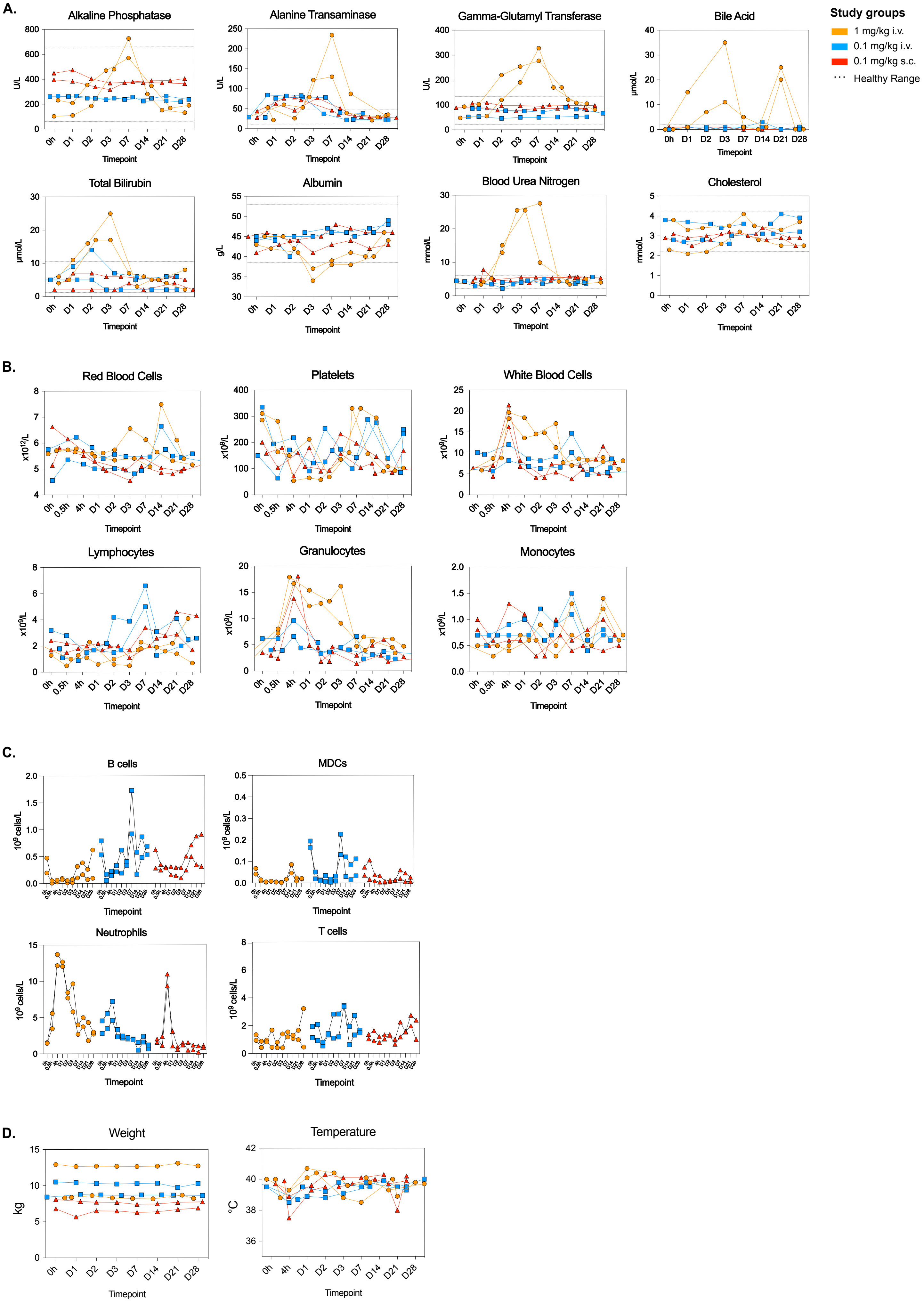

**Supplementary Figure 3**

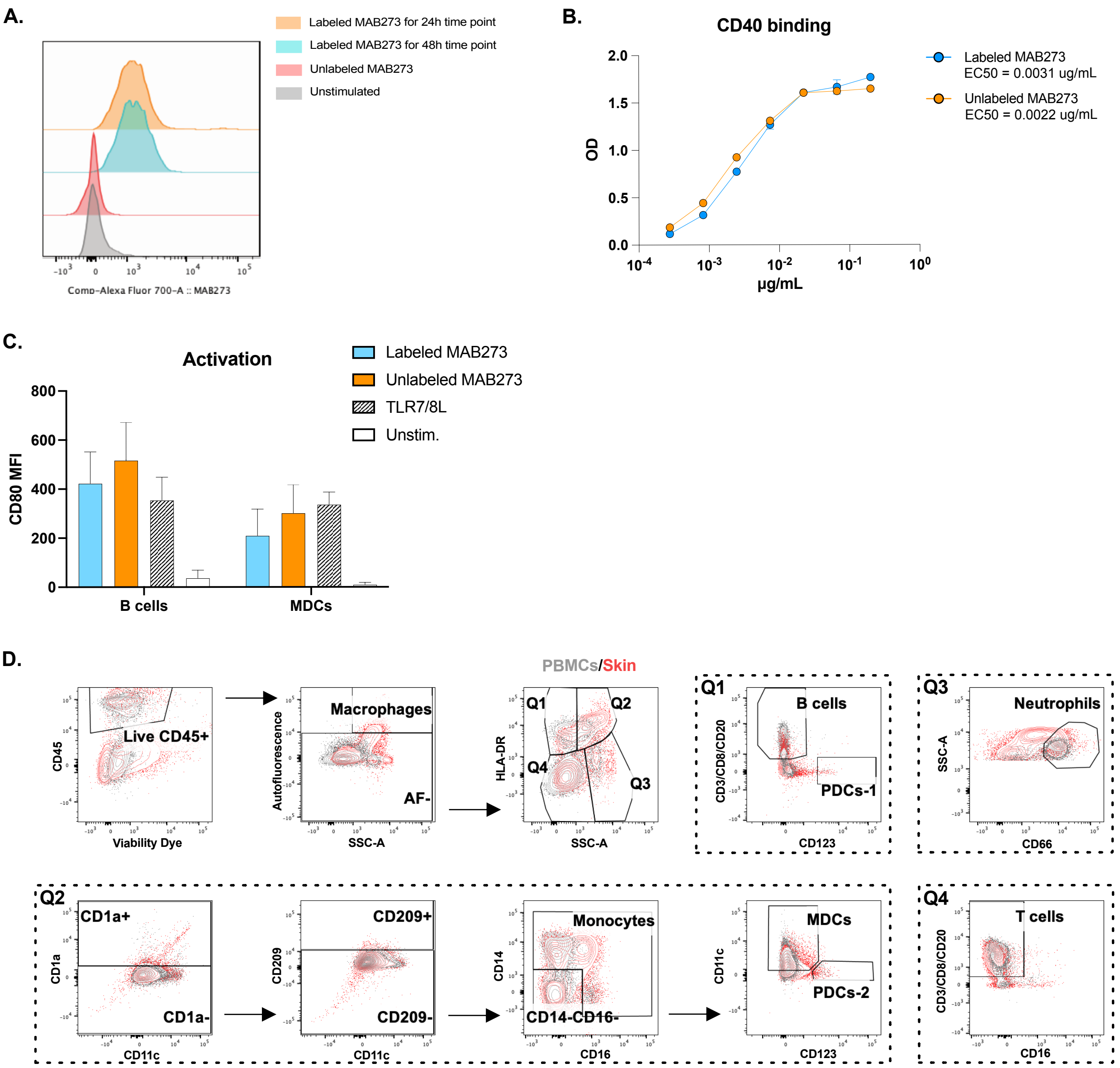

**Supplementary Figure 4**

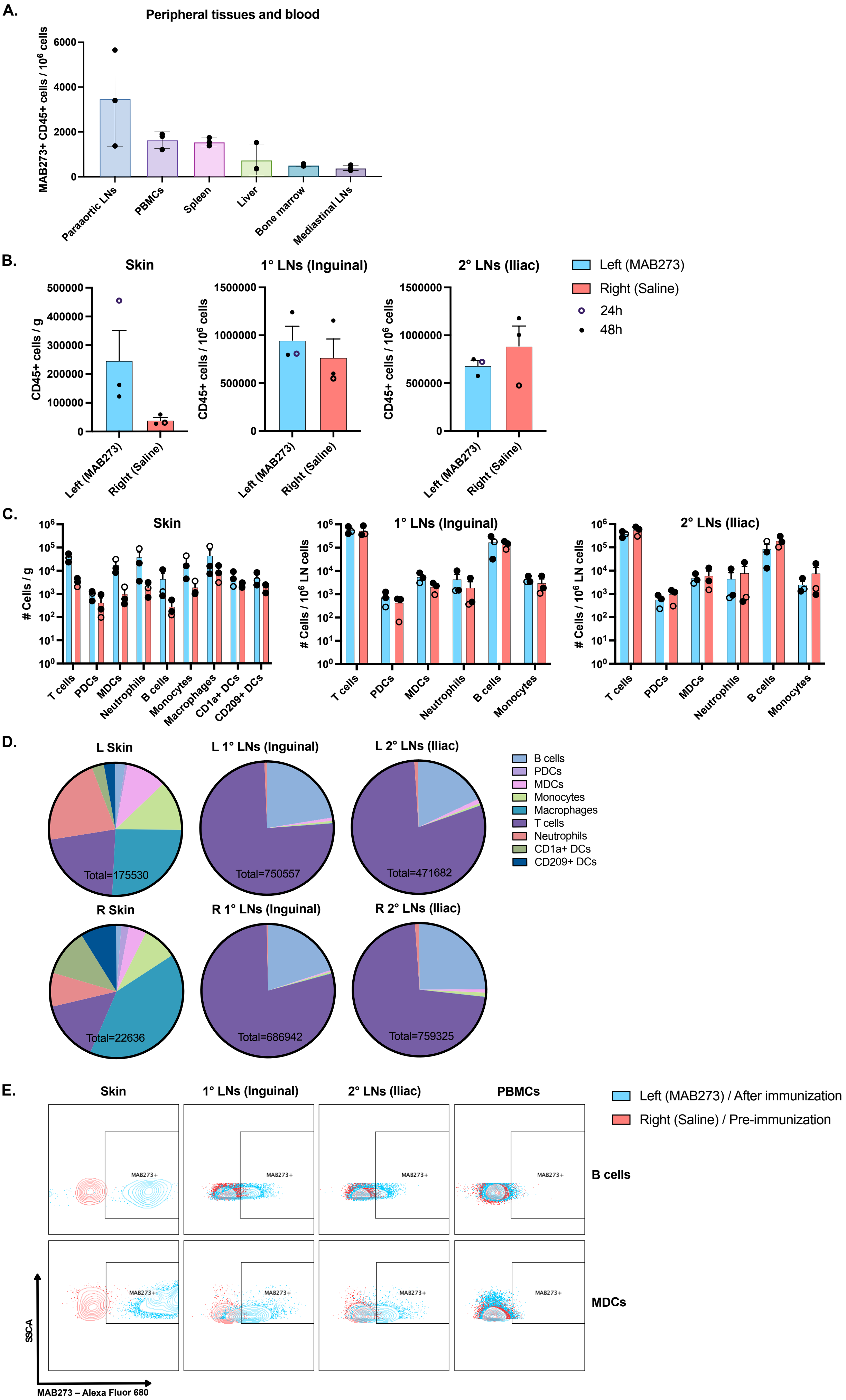

Supplementary Figure 5

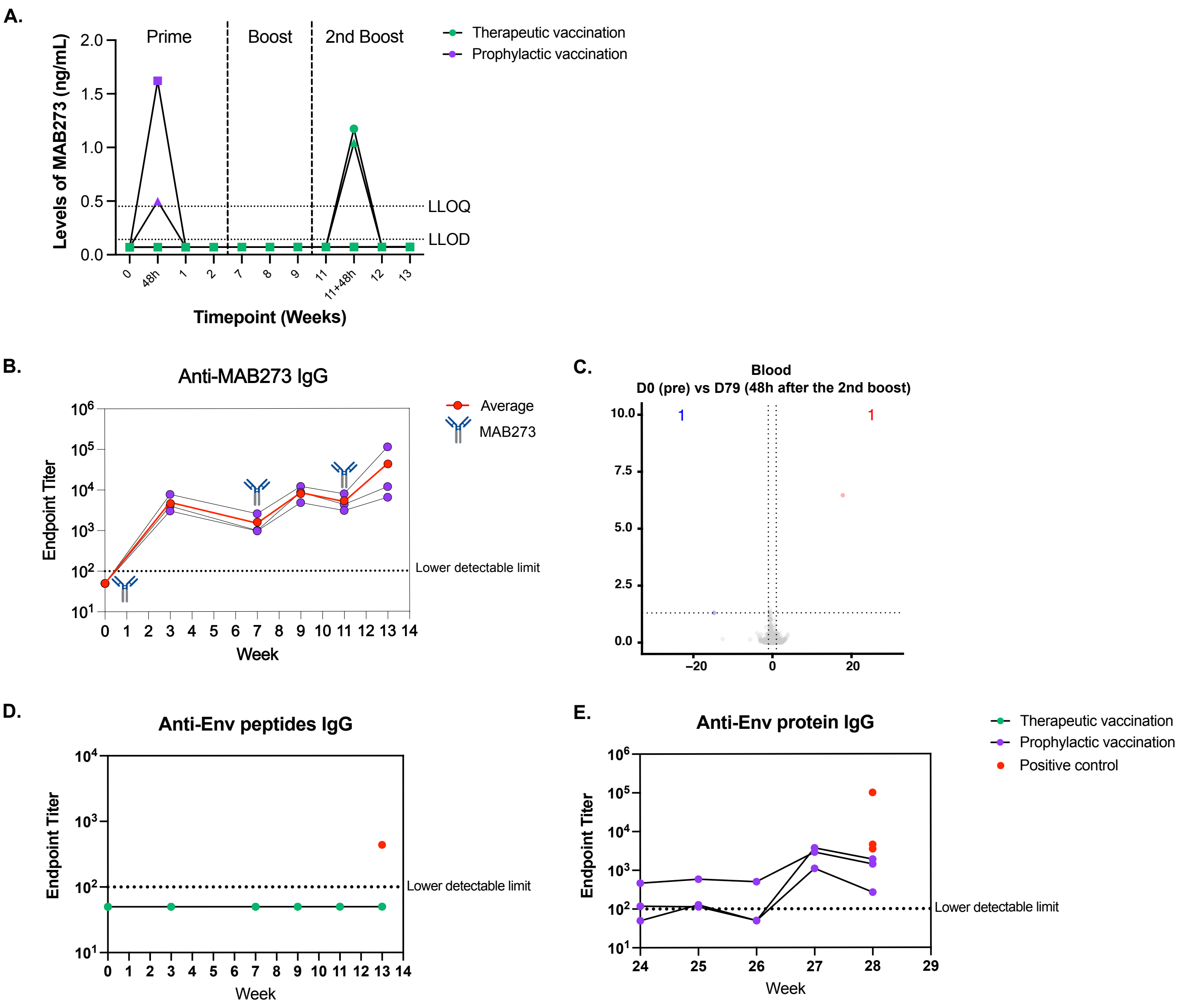

**Supplementary Table 1**

| Antibody | Fluorochrome | Clone | Company | Antibody | Fluorochrome | Clone | Company |
|---|---|---|---|---|---|---|---|
| Innate phenotyping | | | | Antigen-specific T cells surface | | | |
| CD40 | FITC | 5C3 | Biolegend | OX40 | BV510 | L106 | BD |
| NKg2a | PE | Z199 | Beckman | CCR7 | BV786 | G043H7 | Biolegend |
| CD80 | BV421 | L307.4 | BD | CD103 | FITC | 2G5 | Beckman |
| CCR7 | PE-Dazzle 594 | G043H7 | Biolegend | CD8a | BV711 | RPA-T8 | Biolegend |
| CD123 | PerCp-Cy5.5 | 7G3 | BD | CD4 | PE-Cy5.5 | S3.5 | Invitrogen |
| CD3 | APC-Cy7 | SP34-2 | BD | CD45RA | BV650 | 5H9 | BD |
| CD66 | APC | TET2 | Miltenyi | Antigen-specific T cells intracellular | | | |
| CD70 | BV786 | Ki-24 | BD | 4-1BB | APC | 4B4-1 | BD |
| HLA-DR | BV650 | L243 | Biolegend | IL-2 | PE | MQ1-17H12 | BD |
| CD11c | PE-Cy7 | 3.9 | Biolegend | CD69 | ECD | TP1.55.3 | Beckman |
| CD16 | AF700 | 3G8 | BD | CD3 | APC-Cy7 | SP34.2 | BD |
| CD20 | BV605 | 2H7 | Biolegend | IFNg | AF700 | B27 | Biolegend |
| CD14 | BV510 | M5E2 | Biolegend | B cell proliferation | | | |
| Labeled antibody tracking | | | | HLA-DR | PE-Cy5.5 | Tu36 | ThermoFisher |
| CD1a | PE | SK9 | BD | CD3 | APC-Cy7 | SP34-2 | BD |
| CD209 | PerCP-Cy5.5 | DCN46 | BD | CD20 | BV605 | 2H7 | Biolegend |
| CD40 | FITC | 5C3 | Biolegend | CD40 | FITC | 5C3 | Biolegend |
| CD11c | PE-Cy7 | 3.9 | Biolegend | CD40L competition | | | |
| CD14 | BV711 | M5E2 | Biolegend | Streptavidin | PE | | Biolegend |
| CD66 | APC | TET2 | Miltenyi | CD3 | APC-Cy7 | SP34-2 | BD |
| CD45 | BV605 | D058-1283 | BD | HLA-DR | PE-Cy5.5 | Tu36 | ThermoFisher |
| CCR7 | PE-Dazzle 594 | G043H7 | Biolegend | CD11c | PE-Cy7 | 3.9 | Biolegend |
| CD3 | APC-Cy7 | SP34-2 | BD | CD20 | BV605 | 2H7 | Biolegend |
| CD8 | APC-Cy7 | RPA-T8 | BD | CD14 | BV510 | M5E2 | Biolegend |
| CD20 | APC-Cy7 | L27 | BD | CD16 | BV421 | 3G8 | Biolegend |
| HLA-DR | PE-Cy5.5 | Tu36 | ThermoFisher | | | | |
| CD123 | BV510 | 6H6 | Biolegend | | | | |
| CD80 | BV650 | L307.4 | BD | | | | |
| CD16 | BV421 | 3G8 | Biolegend | | | | |
